# An ancient X chromosomal region harbours three genes potentially controlling sex determination in *Cannabis sativa*

**DOI:** 10.1101/2025.07.03.663031

**Authors:** Matteo Toscani, Afsheen Malik, Ainhoa Riera-Begue, Caroline Dowling, Quentin Rougemont, Ricardo C Rodríguez de la Vega, Tatiana Giraud, Susanne Schilling, Rainer Melzer

## Abstract

- Sex determination mechanisms in dioecious plants remain poorly understood yet offer an excellent model system to study genetic changes underlying morphological evolution.
- We investigated the genetic basis of sex determination in *Cannabis sativa*, combining QTL mapping in a segregating population, comparative transcriptomics between monoecious and dioecious cultivars, and a genomic analysis of X-Y chromosome divergence.
- QTL mapping identified *Monoecy1*, a locus on the X chromosome putatively controlling the monoecy-dioecy trait. This locus resides in the most ancient and diverged region of the sex chromosomes and contains three genes within 60,000 bp (*CsREM16*, *lncREM16*, and *CsKAN4*) with distinct sex-specific and monoecy-specific expression patterns.
- *Monoecy1* harbours genes for male-female as well as monoecious-dioecious sex determination. We propose that the combinatorial interaction of *CSREM16*, *lncREM16* and *CsKAN4* provides a unifying genetic framework for understanding male-female and monoecious-dioecious sex determination in *Cannabis sativa*.

## Introduction

The evolution of sex chromosomes remains a major evolutionary puzzle because it involves the emergence of highly differentiated chromosomes from initially homologous autosomes through recombination suppression, a process that is generally disadvantageous in the long term, notably due to reduced selection efficiency and gene loss. The current paradigm suggests that flowering plant sex determination evolved from hermaphroditic ancestors through sequential mutations causing male and female sterility, localized on proto-sex chromosomes (Charlesworth, 2013). In the chromosomal regions harboring the sterility loci, recombination ceases, to avoid producing infertile offspring that would have both sterility alleles. The non-recombining region then often expands over time along the sex chromosomes, generating evolutionary strata of differentiation between sex chromosomes (Bergero & Charlesworth, 2009). Indeed, sequence differences accumulate between sex chromosomes as a function of time since recombination suppression. Furthermore, the recombination suppression leads to less efficient selection, and therefore often degeneration, including gene losses (Bachtrog, 2013).

The size of non-recombining regions varies considerably among species. In asparagus, these regions are relatively small, containing only a very few genes (Harkess *et al*., 2017), facilitating the identification of candidate sex-determining genes. By contrast, *Cannabis sativa* (hemp) features a vast non-recombining region spanning approximately 55Mb (Lynch *et al*., 2025), encompassing the majority of the sex chromosomes and thousands of genes, complicating the identification of sex-determining candidates.

*Cannabis* is a multipurpose crop with applications in the textile industry due to its fiber production, food industry with hemp seeds and hemp seed oil, and medicinal compounds present in its inflorescences such as CBD (Andre *et al*., 2016; Adhikary *et al*., 2021; Schilling *et al*., 2021). Unlike the vast majority of flowering plants that have bisexual flowers with male and female reproductive organs in one flower, *Cannabis* is mainly a dioecious crop, with male and female flowers appearing on separate individuals. However, monoecious hemp cultivars with male and female flowers on the same individual do exist as well. The monoecy trait is agriculturally significant, as monoecious cultivars are preferred for fiber production due to their uniformity because of the absence of male plants that flower and senesce earlier than female plants in dioecious cultivars (Pancaldi *et al*., 2025).

In *Cannabis* the sex is primarily determined by sex chromosomes: male plants possess heterogametic XY chromosomes, while female as well as monoecious plants possess homogametic XX sex chromosomes (Divashuk *et al*., 2014; Faux *et al*., 2014; Riera-Begue *et al*., 2025). However, considerably plasticity exists in the system, the genetic sex can be overridden by environmental cues and hormone treatments and monoecious plants with an XY chromosome set have been identified (Lubell & Brand, 2018; Garcia-de Heer *et al*., 2024; Timoteo Junior & Oswald, 2024; Monthony *et al*., 2026).

In recent years several groups have attempted to identify sex determination genes in *Cannabis*. Those endeavours have led to several candidate genes: Akagi et al. (2025) proposed an *ETR1-like* ethylene receptor gene (*CsETR1*), Carey et al. (2025) proposed a homolog of an aminocyclopropane-1-carboxylate synthase (*CsACS3*) involved in ethylene biosynthesis and Shi et al. (2025) proposed a transcription factor gene belonging to the reproductive meristem gene family (*CsREM16*). All those sex determination candidate genes have in common that they are located on the X chromosome, and an X chromosome-autosome balance system has been suggested as the predominant sex determination system (Akagi *et al*., 2025; Carey *et al*., 2025).

Despite the clear agricultural and theoretical interest in understanding monoecy in hemp, very little is known about its genetic basis. Previous studies found QTLs for monoecy on unplaced scaffolds (Petit *et al*., 2020). A more recent study suggested *CsACS3* as a candidate gene also for monoecy vs dioecy (Carey *et al*., 2025)..

Moreover, despite our previous transcriptomics analysis identifying candidate genes responsible for male/female sex determination (Shi *et al*., 2025), and transcriptomics analysis on hormone-induced male flower development in female plants (Adal *et al*., 2021; Garcia-de Heer *et al*., 2025), comprehensive studies comparing gene expression between female and monoecious hemp cultivars remain scarce. Yet, such studies have been successfully used in other species to pinpoint genes related to the monoecy trait (Zhao *et al*., 2025). Additionally, gene mapping has proven effective in identifying candidate loci related to monoecy in species such as spinach and *Salix purpurea*, which exhibit both dioecious and monoecious lines (Hyden *et al*., 2023; Yamano *et al*., 2024).

Hemp, displaying both monoecy and dioecy, is an ideal candidate to identify their genetic basis and could further advance the more general question of plant sex determination, with sex-determining genes currently identified only in a handful of dioecious and monoecious plant species (Marais *et al*., 2025). Notably, the Cannabaceae hemp and hop (*Humulus lupulus*) possess some of the oldest known pair of sex chromosomes in flowering plants, evolved before the divergence of the two species around 28 million years ago (Prentout *et al*., 2021). Thus, elucidating the sex determination mechanism and the genetic network underlying monoecy may offer critical insights into the evolution of plant sex chromosomes. To pinpoint the genetic basis of sex determination and monoecy in *Cannabis*, we employed three complementary methods: (1) QTL mapping in a segregating population, (2) comparative gene expression analyses between monoecious and dioecious cultivars (and their F_2_progeny), and (3) genomic comparison between the X and Y chromosomes, to assess their degree of differentiation, the possible presence of evolutionary strata and of gene losses.

We show here that the X chromosome underpins not only male-female sex determination but also the monoecy trait. Our methods consistently converge at the most diverged region of the X chromosome, containing both male-female sex determination candidates and the monoecy locus, as central to these processes.

These findings support a model in which a mutated female (XX) genotype, through disruption of the femaleness-determining pathway, gained the capacity to produce male flowers in monoecious cultivars. This study thus provides a unifying genetic framework for understanding the evolution and regulation of sex in *Cannabis sativa*, with implications for breeding programs and for understanding how sex determination pathways evolve on plant sex chromosomes.

## Materials and Methods

### Plant Material and Growth Conditions

The ‘Felina 32’x’FINOLA’ mapping population was described previously (Dowling *et al*., 2024). Briefly, an individual of the dioecious ‘FINOLA’ cultivar was used as pollen donor and the monoecious ‘Felina 32’ cultivar as pollen acceptor. The F_2_ mapping population was generated by crossing a single male F_1_ as a pollinator with four F_1_ female plants. The F_2_ population was grown in a glasshouse under natural light conditions in Dublin, Ireland from June to September 2020. Plants were phenotyped once per week during the observation period and analysed for the presence of male and female flowers. The individuals with both male and female flowers were phenotyped as monoecious while plants with entirely male and female flowers were marked as male and female, respectively.

Leaf tissue was harvested from the F_2_ population and stored at −80°C until further processed. The DNA from all F_2_ individuals, the F_1_ individuals, and the parents was extracted using DNeasy® Plant Mini Kit (Qiagen GmbH, Germany) as per manufacturer’s instructions. The quality and quantity of DNA was checked by agarose gel electrophoresis with a NanoDrop spectrophotometer (ND-1000). The samples were prepared to a final concentration of ≥ 35 ng/µl and sent to BMKGene (Beijing, China) for SLAF-Seq (Specific Locus Amplified Fragment Sequencing).

The F_4_ population was grown under glasshouse conditions from June to September 2024 under glasshouse conditions in Dublin, Ireland. The floral meristem and axillary inflorescence meristem were sampled for RNA extraction. The RNA extraction was carried out using RNeasy® Plant Mini Kit (Qiagen GmbH, Germany) as per manufacturer’s instructions. The RNA concentration was determined using Nanodrop spectrophotometer (ND-1000), while RNA integrity was determined using Bioanalyzer (Agilent). The samples with RIN > 5 were further processed for RNA Sequencing, which was carried out by Novogene (Cambridge, UK).

### SLAF-seq and quality control

A reduced representation genome sequencing approach, specific locus amplified fragment sequencing (SLAF-seq) (Sun *et al*., 2013), was used to sequence the F_2_ population. SALF-seq was carried out by BMKgene. The service provider (BMKGene) conducted a simulation experiment for selecting the most appropriate restriction enzymes against the hemp reference genome assembly (GF_029168945.1_ASM2916894v1) followed by their testing of selected samples in a pilot experiment. From the simulation and pilot experiment results, the restriction enzyme set RsaI and HaeIII was selected as the most appropriate set generating a substantial number of fragments (SLAF-tags) that are randomly and uniformly distributed across the genome and avoid repetitive regions. SLAF-tags were about 364 - 444 bp in size.

On average 12640 SLAF tags were produced per sample with a mean sequencing depth of 20X. Libraries generated from SLAF-tags after a series of steps were processed on the Illumina platform and about 51.60 Mb reads were generated. The quality of SLAF-seq fastq files was checked using fastQC and summarised using MultiQC softwares (Ewels *et al*., 2016).

All reads exceeded quality 20 and consequently no filtering was applied at this stage. The adapter sequences were trimmed by the BMKgene sequencing company and consequently no further trimming was performed.

### Mapping and genotyping

After quality control, the paired-end reads from female and monoecious samples were mapped against the Pink pepper reference genome (GCF_029168945.1_ASM2916894v1) using BWA-mem2 (Vasimuddin *et al*., 2019). The reads from male samples were mapped against the reference genome to which the non-recombining region of the Y chromosome from the ‘Kompolti’ assembly (Lynch *et al*., 2025) (*KOMPb*) was appended, to avoid reads from the Y chromosome getting mapped to similar regions in the X chromosome and falsely genotyping heterozygous SNPs in male X chromosome positions. We then measured for each sample the proportion of reads mapping to the Y chromosome.

The resulting BAM files were filtered to keep reads of mapping quality MAPQ > 20. The analysis was then repeated for the reads that mapped against the chromosome X with the higher threshold filtering MAPQ > 35. Due to SLAF-seq reads being repetitive by design, duplicate reads were not removed. The genotyping was then done using GATK 4.6.0 functions HaplotypeCaller and GenotypeGVCFs (DePristo *et al*., 2011).

### QTL analysis

The resulting VCF file from GenotypeGVCFs, containing SNPs from the mapping population, was filtered using VCFtools 0.1.17 (Danecek *et al*., 2011) to keep biallelic markers that were genotyped in at least 80 % of the samples, with quality greater than 30 and with depth of at least 10, with the following parameters: --min-alleles 2 --max-alleles 2 --max-missing 0.8 -- minQ 30 --minGQ 30 --minDP 10.

The genotypes from the filtered VCF file were then imported in R using the package onemap (Margarido *et al*., 2007), which allowed also to convert the VCF file into a genotype table coded by the parental origin: markers from F_2_ samples which were homozygous for the ‘FINOLA’ parent were coded as AA, markers homozygous for the monoecious ‘Felina 32’ parent were coded as BB, heterozygous markers from both parents were coded as AB. To this genotype table, the phenotype data was then manually added, with the plant sex encoded as 1 for female and 0 for monoecious, in columns as required by R/qtl (Broman *et al*., 2003) cross format, and the cross object was imported with the read.cross() function. For the QTL analysis, female and monoecious plants were used, as male plants inherit the whole Y chromosome. Markers from the autosomes which show significantly distorted segregation (P.value < 0.05 after applying a Bonferroni correction for multiple testing) from the expected AA:AB:BB 1:2:1 ratio were removed. For each chromosome, markers were reordered based on their recombination frequency using the R/qtl orderMarkers() function. The genetic map in centimorgans is then estimated with the est.map() function, with parameters map.function = “kosambi”,error.prob = 0.01, with average and median chromosomal lengths of 105.5 cM and 97.67 cM, respectively. The average marker spacing across chromosomes was <1 cM. Low recombination fractions for markers on the same chromosomes, together with high Logarithm of the Odds (LOD) scores, were used as indicators to confirm that markers assigned to the same linkage groups were closely linked, as expected for markers within the same chromosome, to evaluate the accuracy of our genetic map and its concordance with the reference genome.

For each phenotypic trait the QTL LOD is calculated with the scanone function and method “mr-argmax”, and the 5 % significance threshold is generated by executing 10,000 permutations. To refine the significant QTL to a more precise genomic interval, we additionally applied the composite interval mapping method using the cim() function in the R/qtl package, using the same genetic map and phenotypic dataframe, improving resolution by fitting multiple marker covariates outside the testing interval to control for genetic background noise, thereby increasing the power to detect QTL, and potentially distinguish closely linked loci. In our case, we used the default model settings, with the default window of 10 cM, without specifying additional covariates, relying on the automatic forward selection procedure from the r/qtl package to select cofactors from the genome scan. The automatic cim forward selection procedure selected and used for the analysis 3 marker covariates: X_81454470, 4_42194662 and 1_43452032.

The same QTL analysis with equal parameters was then repeated after remapping the reads against both haplotypes SAN2a and SAN2b of the phased ‘Santhica’ assembly (Lynch et al., 2025).

### Gene loss ratio estimate

The degree of degeneration of different segments of the Y chromosome was estimated using the gene loss ratio as a proxy for divergence with the X chromosome. The X chromosome was divided into 10 bins of maximum 10 million nucleotides. From each bin all the proteins encoded by genes in that chromosomal section were BLASTed against the genomic sequences of the Y chromosome assemblies of 3 different cultivars (Lynch *et al*., 2025): Kompolti (KOMPa), BooneCounty (BCMa), and a landrace cultivar from Germany (GRMa), using the tblastn algorithm. If no match with e-value < 0.05 was found in any of the assemblies, a gene was considered to have been lost on the Y chromosome. The percentage of X chromosomal genes with no counterpart on the Y chromosome was calculated for each bin and plotted.

We consider this approach a conservative estimate of gene loss due to the fact that a gene present on the X chromosome belonging to a gene family can have a match on the Y chromosome if a close member of this family is encoded on the Y chromosome, hence the “true” gene loss ratio segment is likely to be higher than the numbers estimated using this approach.

For each X chromosome section bin, additionally, the percentage of female biased genes was calculated based on previously published transcriptomics analyses (Shi *et al*., 2025). Genes that were overexpressed in females at either the vegetative or flowering stage were counted as female-biased.

### Differentiation between sex chromosomes: synonymous divergence (d_S_) and evolutionary strata

From the available phased Kompolti assembly (KOMPa, KOMPb) (Lynch *et al*., 2025), the haplotypes containing X and Y chromosomes were selected and synonymous divergence (d_S_) values for single-copy orthologs calculated and the detection of evolutionary strata conducted through the EASYstrata pipeline (Rougemont *et al*., 2025). The pipeline identifies single-copy orthologs between X and Y chromosomes using OrthoFinder (Emms & Kelly, 2019). Then it aligns coding sequences for each orthologous pair accounting for frameshifts and stop codons. In the next step, per-gene synonymous divergence (d_S_) and non-synonymous divergence (d_N_) are calculated using the codeml program implemented in PAML (Yang, 2007). The resulting d_S_ values are plotted along the X chromosome gene order to visualize patterns of divergence along an ancestral-like gene order. Evolutionary strata are then inferred using Bayesian changepoint analysis implemented in the MCP R package (Lindeløv, 2020), which detects discontinuous changes in mean d_S_ values along the chromosome, for different change-point values.

We then aimed to identify the evolutionary strata older than the *Cannabis*-*Humulus* speciation. When recombination suppression precedes speciation, substitutions remain associated with the same sex chromosome through speciation. Therefore, genes in those strata would be predicted to contain trans-specific polymorphisms, i.e. in a phylogenetic tree X chromosomal genes from *Cannabis* would be more closely related to X chromosomal genes from *Humulus* than to Y chromosomal genes from *Cannabis*. We identified a subset of 313 genes that were single-copy orthologs between X and Y chromosomes in both *Cannabis* and *Humulus*. We then constructed single-gene genealogies to identify those with trans-specific polymorphisms based on the tree topology.

### Transcriptomics analysis

RNA-seq reads were generated previously (Dowling *et al*., 2024) from leaves samples from monoecious ‘Felina 32’ and dioecious ‘FINOLA’ females. They were now mapped against the transcriptome of the reference genome ‘Pink pepper’ using Salmon, gene expression was quantified and a comparative analysis of differential gene expression was performed using the R Deseq2 package (Love *et al*., 2014). The same comparison was performed between 3 F_2_ female and 3 F_2_ monoecious individuals with leaf samples that were obtained from the mapping population. An F_3_ and F_4_ population was generated from the F_2_ population by random intercrossing. From the F_4_ population 8 monoecious and 8 female dioecious samples from floral meristem and axillary inflorescence meristem were taken and analyzed with the same aforementioned pipeline. In DESeq2, adjusted p-values (padj) are set to NA for genes with extremely low expression, zero variance across samples, or in cases where the statistical model fails to converge, often due to a small sample size causing high dispersion. Our F₂ transcriptome dataset, comprising three biological replicates per group, may contribute to increased variability and dispersion, leading to a substantial number of genes with NA padj, including some with extremely low unadjusted p-values. To identify genes consistently downregulated in monoecious plants across generations, we applied a filtering threshold of p < 0.05 and |log₂ fold change| > 1, to retain differentially expressed genes regardless of padj availability.

We analyzed the expression of three candidate sex-determination genes (*CsKAN4*, *CsREM16*, *lncREM16*) in the 64 male, female, and monoecious samples from the cultivars ‘Felina 32’ and ‘FINOLA’s part of the previously published experiment mentioned above (Dowling *et al*., 2024) and publicly available under BioProject PRJNA956491.

Expression of homologs of candidate sex determining genes *CsKAN4* and *CsREM16* was evaluated in *Humulus* using published RNA-seq data from five male and five female hop plants available under BioProject PRJNA694508. Homologs were identified through a BLAST search, the gene expression was quantified though Salmon and the statistical significance evaluated through the R Deseq2 package.

To compare expression patterns of our candidate sex-determining genes with previously proposed candidates, we used RNA-Seq data from our earlier experiment (Shi *et al.,* 2025) (BioProject accession PRJNA1126191). Briefly, directional RNA-Seq (PE150, Novogene, UK) was performed on 20 samples comprising four biological replicates each of two-leaf stage male (L2M) and female (L2F), four-leaf stage male (L4M) and female (L4F), and nine-leaf stage female (L9F) plants. Moreover, to investigate whether the expression of candidate sex-determining genes is affected by phytohormones treatment, we analysed published RNA-seq data from *Cannabis sativa* male (XY), female (XX), and silver thiosulfate-induced male (XX) flowers (Adal et al., 2021; BioProject PRJNA669389). Expression was quantified using Salmon, and differential expression analysis was performed using DESeq2, with sex (male, female, silver-induced male) as the experimental factor. Pairwise contrasts between all three groups were computed and p-values were adjusted using the Benjamini-Hochberg method.

### *CsKAN4* expression analysis

‘FINOLA’ and ‘Felina 32’ seeds were sown on a mixture of compost, John Innes no2, vermiculite and perlite (2:1:1:1). After two days of incubation in darkness to promote germination, lights (red and blue LEDs) were turned on with 18:6 photoperiod during 2 weeks for vegetative growth, and then photoperiod changed to 14:10 to induce flowering.

To study the expression of *CsKAN4* in ‘FINOLA’ and Felina32, apical meristem samples were taken at 1st leaf, 2nd leaf and 4th leaf stage. In the case of the ‘’FINOLA’, from a part of the sample DNA was isolated and genotyping was performed using the *CsPDS5*-CAPS protocol described in Riera-Begue et al. (2025) to determine if the plants were male or female.

From the remaining sample RNA isolation was carried out using the RNeasy® Plant Mini Kit (Qiagen, Germany) according to the manufacturer’s instructions.

For qPCR, 200 ng of the isolated RNA was used to synthesize cDNA following the SuperScript™ IV Reverse Transcriptase (Invitrogen™) protocol as per supplier’s instructions. The primer sequences used for the qPCR are the following:

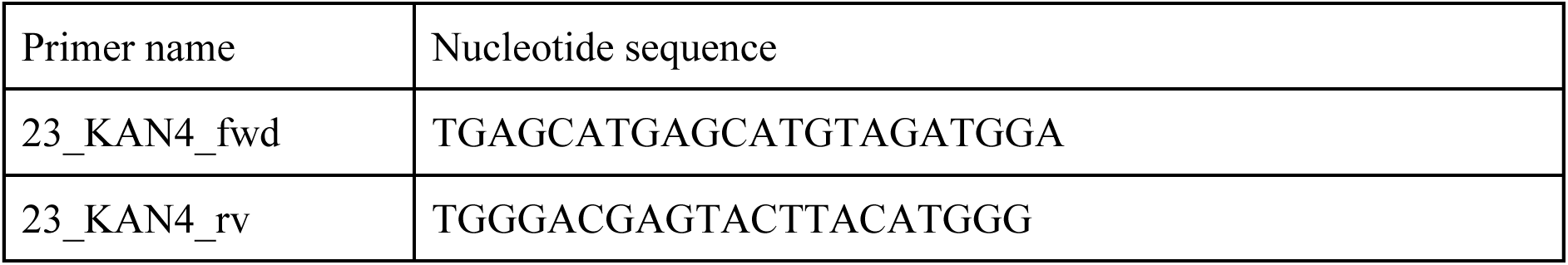

qPCRs were conducted in a final volume of 10 μL with: 5 μL Fast SYBR™ Green Master Mix (ThermoFisher Scientific), 0.2 μL of 23_KAN4_fwd (10 μM), 0.2 μL of 23_KAN4_rv (10 μM), 0.6 μL DEPC water and 4 μL of the cDNA template (2 ng/μL).

The QuantStudio™ 5 Real-Time PCR System (ThermoFisher Scientific) was used for the RT-qPCR. After an initial hold stage at 95 °C for 20 s, 40 cycles at 95 °C for 10 s, at 58 °C annealing temperature for 30 s were performed before a melt curve stage which consisted in 95 °C for 15 s, 60 °C for 80 s and 95 °C for 15s.

Expression levels were normalized to *CsPP2A* and *CsUBQ* as housekeeping genes (Guo et al., 2018).

## Results

### Investigation of evolutionary strata and shared recombination suppression between Cannabis *and* Humulus

To identify the region of the sex chromosomes likely to harbor the sex determining genes, we searched for the oldest area of recombination suppression between the X and Y chromosomes, as this region would have been the first to cease recombining and would therefore show the highest sequence divergence. To search for evolutionary strata, we plotted synonymous divergence (d_S_) for single-copy X-Y orthologs in the *Cannabis* cultivar ‘Kompolti’ (Fig. 1a,b, Supplementary Table S1) and in *Humulus* along the X chromosome (Fig. 1c,d), as d_S_ increases with time since recombination suppression. For both species, the changepoint analysis inferred a single evolutionary stratum for the most supported model (Supplementary Fig. S1). A less supported model with two changepoints inferred a small younger evolutionary stratum expanding the non-recombining region towards the pseudo-autosomal region.

**Figure 1.**
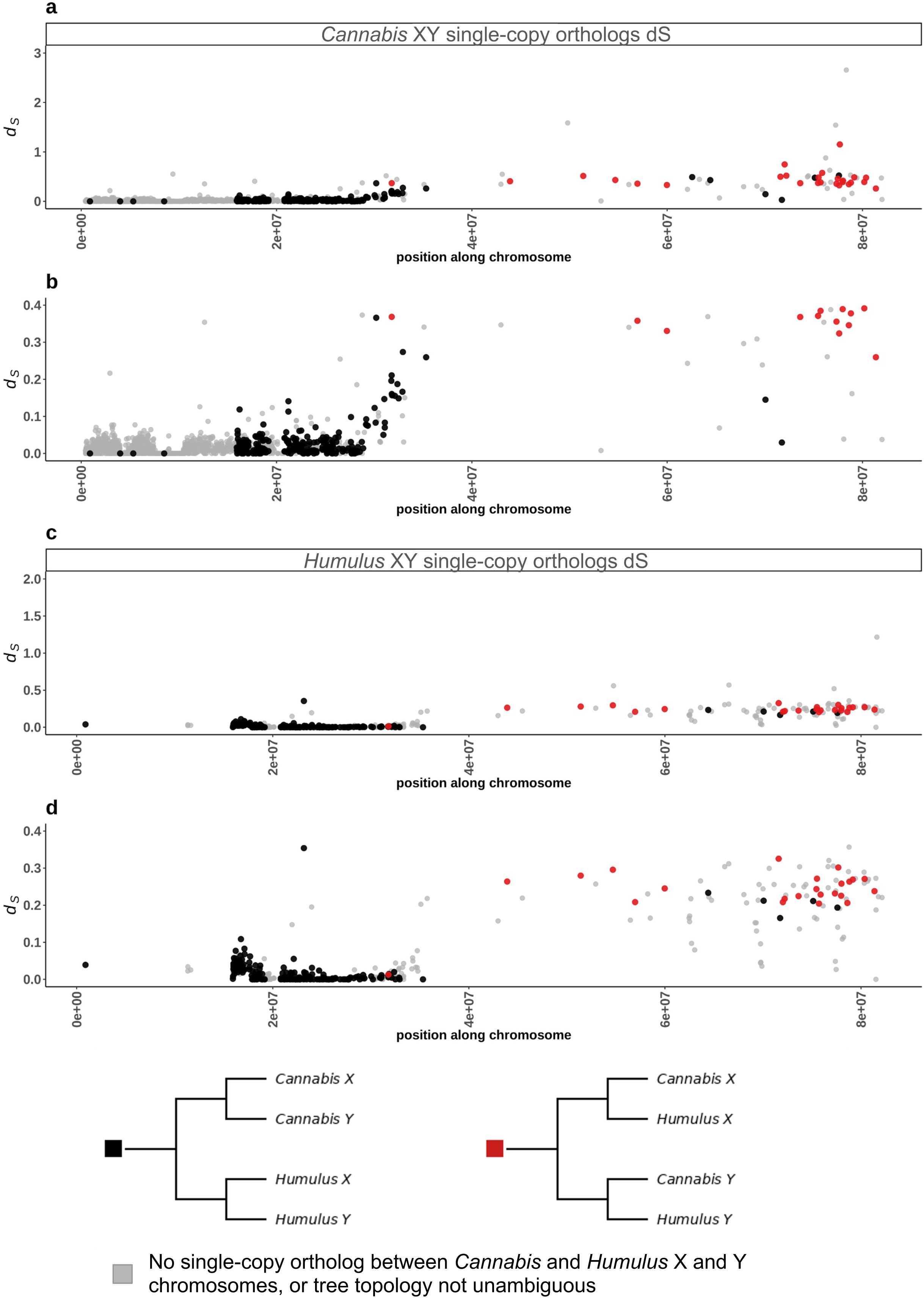
Synonymous divergence (d_S_) and trans-specific polymorphisms analysis of *Cannabis* and *Humulus* X-Y single copy orthologs. **a)** Synonymous divergence substitution rates (d_S_) between *Cannabis sativa* X-Y single-copy orthologs plotted against their chromosomal position in the X (Mb). The y-axis is truncated at d_S_ = 3 to allow visualization of the majority of gene pairs while excluding extreme outliers. **b)** *Cannabis* X-Y d_S_ values plotted by gene rank order along the reference chromosome. The y-axis is truncated at d_S_ = 0.4 to improve resolution of functionally constrained genes and exclude possible pseudogenes. **c)** Synonymous divergence substitution rates (d_S_) between *Humulus lupulus* X-Y single-copy orthologs plotted against their chromosomal position in *Cannabis sativa* X (Mb), with y-axis truncated at d_S_ = 3. **d)** *Humulus* X-Y d_S_ values plotted by gene rank order along the reference chromosome, along the x-axis, with y-axis truncated at d_S_ = 0.4. In all panels, genes with trans-specific polymorphisms, presumed to belong to the genomic regions that stopped recombining before speciation, are highlighted in red. Gene ordering is always based on the position of orthologs on the YMv2a assembly chrX (Lynch et al., 2025), which serves here as the “ancestral” reference to maintain a consistent and comparable ordering across the two species.

By anchoring the gene order to the same *Cannabis* physical map, we found that the evolutionary strata inferred in the two species seemed concordant, suggesting that recombination cessation occurred prior to the speciation event separating *Cannabis* from *Humulus*. This was subsequently confirmed though single tree phylogenies identifying genes with trans-specific polymorphisms between *Cannabis* and *Humulus* sex chromosomes (highlighted in red in Fig. 1). The genes with trans-specific polymorphisms were contained in an area spanning from 36 Mb towards the end of the chromosome X, again pointing to the oldest region of recombination suppression being in this area (Supplementary Data S1).

We further identified specific genes exhibiting saturation of synonymous sites (dS>1), indicating multiple synonymous mutations per site on average. The genes displaying the highest divergence, LOC115700360 and LOC115699965, were located at 81.2 Mb and 82.3 Mb on the *Cannabis* X chromosome, respectively.

### Inheritance of monoecy reveals likely X-linked control system

Monoecious and dioecious hemp cultivars are interfertile, which offers the opportunity to analyse the monoecy-dioecy sex determination system genetically using QTL analyses. To elucidate the genetic basis of monoecy, we analyzed segregation patterns across three generations of a previously described cross between the monoecious cultivar ‘Felina 32’ acting as pollen receiver and the dioecious ‘FINOLA’ cultivar acting as pollen donor (Dowling *et al*., ^2^_024_^)^. The F_1_ generation consisted exclusively of dioecious plants (49% male, 44.9% female, with the remaining 6.1% not flowering) with no monoecious individuals among 98 phenotyped plants (Fig. 2a, Supplementary Table S2), demonstrating complete dominance of dioecy.

**Figure 2.**
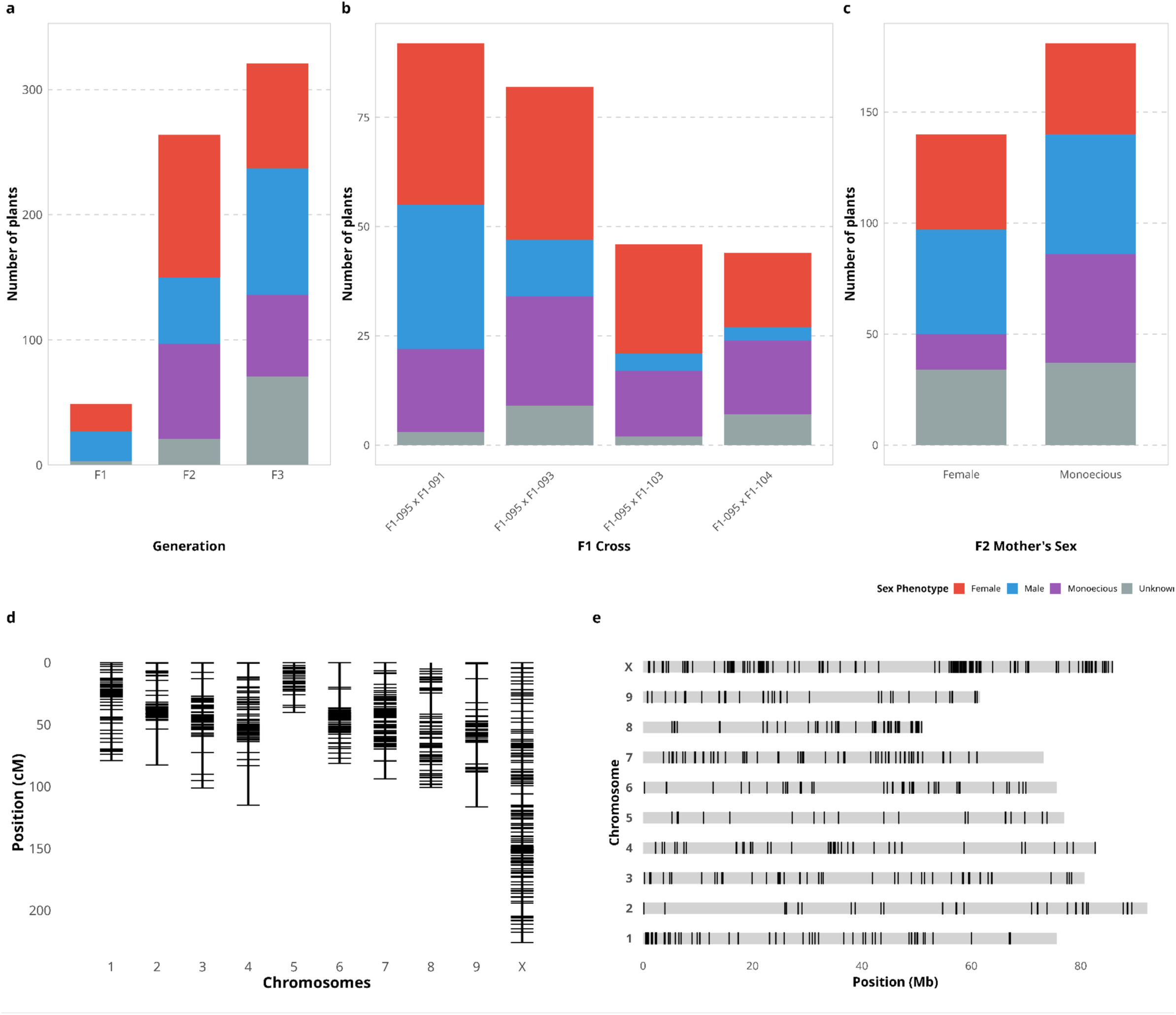
Sex inheritance patterns across generations and genetic mapping in Cannabis sativa. **a)** Sex distribution across three generations of a ‘FINOLA’ (dioecious) × ‘Felina 32’ (monoecious) cross. **b)** Sex distribution of F₂ plants grouped by F₁ mother plant. **c)** Sex distribution of F₃ plants grouped by F₂ maternal parent sex phenotype. **d)** Genetic map constructed from 1,400383 SLAF-seq markers spanning 1,055 cM across 10 chromosomes. **e)** Physical map showing marker distribution along chromosomes in megabases (Mb). Figure 2. QTL mapping and sex chromosome evolution in *Cannabis sativa*.

In the F_2_ generation, monoecy reappeared with complex segregation patterns. Among 242 flowering individuals, we observed 43 % females, 29 % monoecious, and only 20.1 % males, significantly deviating from the expected 50 % male percentage (χ² = 18.3, p < 0.001; Fig. 2a). To produce the F_2_ population, several F_1_ female individuals were pollinated using the same F_1_ male plant. Sex frequencies in the F_2_ progeny varied significantly depending on the identity of the F_1_ mother plant. Despite using the same F_1_ male pollinator, male offspring frequency ranged from 6.8% to 35.9% across different F_1_ mothers (χ² = 23.65, df = 6, p = 0.0006; Fig. 2b), indicating maternal influence on sex inheritance.

252 F_2_ individuals were sequenced using SLAF-seq (see below), and F_1_ plants used for crossing were subjected to whole genome sequencing (Dowling et al., 2024). The correlation between the F_1_ and F_2_ plant sex phenotype and genotype was confirmed by measuring the percentage of sequencing reads mapping to the Y chromosome for each plant, considered here a proxy for the presence of the Y chromosome. The percentage of reads mapped against the Y chromosome was > 6 % in every male sample and ≤ 3 % in female and monoecious plants (Supplementary Table S3), indicating a strong correlation between genetic and phenotypic sex.

Moreover, the sex genotype was confirmed by a previously described PCR assay (Riera-Begue ^e^_t al., 2025_) for 137 of the F_2_ plants, confirming a 100% agreement between the phenotyped and genotyped sex of the tested plants. In particular, phenotyped females and monoecious plants were always genotyped as having XX chromosomes (Riera-Begue et al., 2025).

The F_3_ generation, produced from the F_2_ via random intercrossing, consisted overall of 26 % female, 20 % monoecious, 31 % male individuals and 22 % of plants which did not flower in the observation period. It confirmed a recessive inheritance type of the monoecy trait: monoecious F_2_ mothers produced 34 % monoecious offspring versus only 15.1 % monoecious offspring from female F_2_ mothers, a 2.3-fold difference (χ² = 11.783, df = 2, p = 0.0028; Fig. 2c, Supplementary Fig. S2).

### A high-resolution genetic map identifies the *Monoecy1* locus on the X chromosome

We constructed a high-density genetic map using SLAF-seq analysis of 252 F_2_ individuals. After filtering for quality and segregation distortion, we retained 1400 markers spanning 1,055 centiMorgans across all chromosomes, with an average spacing of <1 cM. The physical map covered approximately 86 Mb of the X chromosome, providing sufficient resolution for QTL mapping (Fig. 2d,e, Supplementary Figs. S3, S4).

QTL analysis using 189 female and monoecious F_2_ plants revealed no significant associations between the monoecy trait and any autosome. However, almost the entire X chromosome showed a strong signal with a peak LOD score of 6.97 at marker X_80375543 (80.38 Mb), exceeding the genome-wide significance threshold of 3.4 (10,000 permutations; Fig. 3a, Supplementary Table S4).

**Figure 3.**
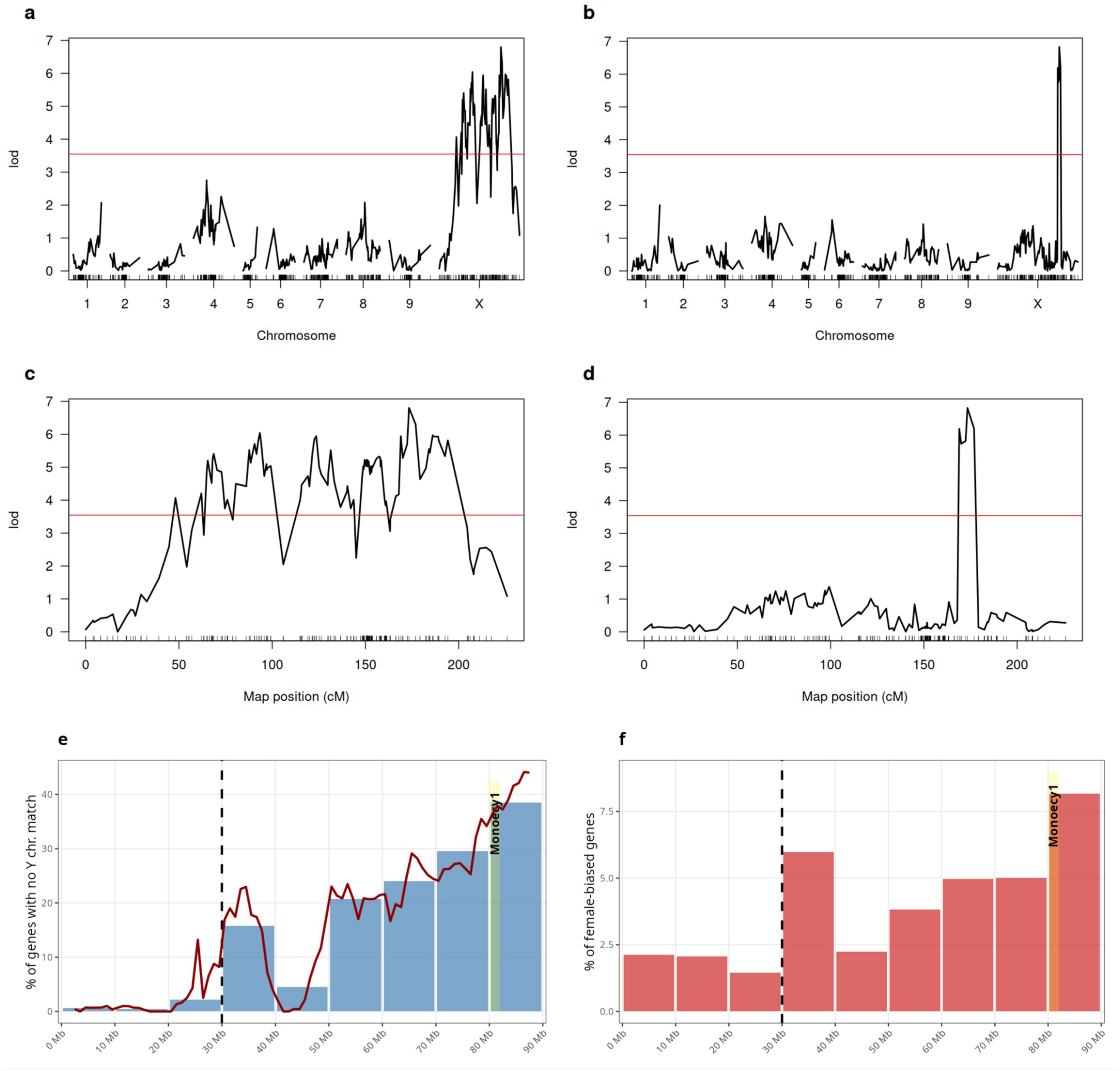
QTL mapping and sex chromosome evolution in *Cannabis*. **a)** Genome-wide QTL scan for monoecy trait using binary model (EM algorithm). The red line indicates the 5% significance threshold (10,000 permutations). **b)** Composite Interval Mapping (CIM) confirming X chromosome QTL. **c)** Detailed X chromosome QTL analysis showing peak LOD score at marker X_80375543. **d)** CIM refinement of *Monoecy1* locus to 80-82 Mb interval. **e)** Percentage of X-chromosomal genes lacking Y chromosome homologs in 10 Mb bins. The dashed line indicates PAR boundary; yellow highlights *Monoecy1*. **f)** Distribution of female-biased genes along the X chromosome. Yellow highlights *Monoecy1*. Female-biased genes are defined as those exhibiting significantly higher expression in females compared to males (adjusted p-value < 0.05, log2 fold change > 1) in at least one of two developmental stage comparisons (vegetative or flowering), as identified in Shi et al. (2025).

As QTL mapping showed a broad peak covering almost the entire X chromosome (Fig. 3a, c), we used Composite Interval Mapping to further refine the signal. This resulted in the identification of a 2 Mb region (80-82 Mb) on the X chromosome (Fig. 3b,d, Supplementary Table S5) strongly associated with monoecy and also peaking at marker X_80375543. This locus, which we designate *Monoecy1*, explains 15% of phenotypic variance for the monoecy/dioecy trait. Notably, *Monoecy1* was located within the area on the X chromosome where the genes displaying the highest d_S_ divergence are contained (Figure 1a).

Further genotype-phenotype association analysis (Supplementary Fig. S5), comparing the genotype at the identified marker X_80375543 (X_803_ hereafter) with the monoecy/dioecy phenotype of the F_2_ plants, provided support for the *Monoecy1* locus (Supplementary Table S6). Individuals homozygous for allele X_803_D_ (which is the parental dioecious ‘FINOLA’ genotype) exhibited an almost exclusively dioecious female phenotype (88 %, although the X_803_D_X_803_D_ genotype was unexpected in F_2_ plants, see Supplementary Text 1, Supplementary Fig. S6, S7). In contrast, genotype X_803_M_, the parental monoecious Felina 32 genotype, was predominantly associated with monoecy (68 % homozygous F_2_ plants being monoecious). Individuals heterozygous for the parental genotypes (X_803_D_X_803_M_) showed an intermediate distribution (62 % female, 38 % monoecious). An analysis of recombination breakpoints confirmed the quantitative nature of *Monoecy1* but did not allow to narrow down the QTL further (Supplementary Figure S8). Repeating the QTL analysis on chromosome X after filtering the reads with a stricter mapping quality threshold of MAPQ > 35 resulted in fewer markers retained but the peak marker remained the same (Supplementary Fig. S9). To avoid the possibility that a genomic region only present on monoecious cultivars was being missed when using the reference ‘Pink Pepper’ genome (Ryu *et al*., 2024) (which is based on a female plant of a dioecious cultivar), we repeated the QTL analysis after mapping the SLAF-seq reads and genotyping against both haplotypes of the monoecious ‘Santhica’ cultivar genome (Lynch *et al*., 2025). The same pattern was obtained (Supplementary Fig. S10), with most of the non-recombining region being associated with the monoecy trait, and the markers with peak LOD significance being X_3257050 and X_3100042, respectively, for the SANa and the SAN2b. By BLASTing the surrounding genomic region, both markers were found to be placed in a region situated in the reference ‘Pink Pepper’ X chromosome at 82.79 Mb.

### Genomic comparisons point to the end of the X chromosome as the most divergent section

To investigate the degeneration of the Y chromosome, as expected following recombination suppression, we estimated the proportion of protein-coding genes that were present on the X chromosome but absent on the Y chromosome. Overall, 418 out of 3,446 (12.1 %) of protein coding genes on chromosome X did not have a counterpart on chromosome Y. The positions of those hemizygous genes were not evenly distributed along the chromosome. The terminal region spanning from 80 Mb to the end of the X chromosome displayed the highest levels of divergence with the Y chromosome, with a conservative estimate of 40 % of X-encoded proteins lacking a Y counterpart (Fig. 3e, Supplementary Table S7). This result is seen in the reference genome but was also confirmed for the alternative ‘Kompolti’ cultivar genome (Supplementary Fig. S11).

Furthermore, there is a clear pattern of the gene loss ratio increasing from one end of the X chromosome to the opposite end, being near zero in the section from 0-30 Mbp. This is in agreement with the position of the previously described pseudo-autosomal region (Prentout et al., 2020) which is still actively recombining between X and Y chromosomes.

An inverse analysis was also performed measuring the genes on the Y chromosome that are not present on the X chromosome. This showed a much lower ratio of Y-exclusive genes (Supplementary Fig. S12), with only around 6% of proteins from the Y exclusive region not found on the X, consistent with the general observation that X chromosomes are typically not actively degenerating like Y chromosomes (Charlesworth, 2013).

Together, our observations may be taken as indication that the sex chromosome evolution in *C. sativa* was initiated in a region towards the end of the X chromosome. This conclusion is further supported by the previous synonymous substitution rate (d_S_) analysis between X and Y chromosome homologs (Prentout et al., 2020, 2021), where a peak in d_S_ is also observed in the terminal section of chromosome X. Interestingly, *Monoecy1*, i.e. the locus controlling monoecy vs. dioecy, is located in the same terminal section of the X chromosome We therefore next analysed data from our previous transcriptomics experiments (Shi *et al*., 2025), in which we compared male with female apical meristem samples at different developmental stages to identify genes with sex-biased expression. When analysing the percentage of X chromosomal genes that showed a female-male biased expression we detected a pattern similar to the gene loss analysis: the greatest percentage of female biased genes was found at the end of the X chromosome, in the region where *Monoecy1* is located (Fig. 3f), with as much as 8 % of the genes in this region being overexpressed in females as compared to males.

### A *KANADI* transcription factor gene is located in *Monoecy1* and differentially expressed between monoecious and female dioecious plants

We next reasoned that candidate genes controlling monoecy might be differentially expressed between monoecious and dioecious cultivars. Initial comparisons between dioecious female ‘FINOLA’ and monoecious ‘Felina 32’ plants revealed 3,420 differentially expressed genes (Fig. 4a, Supplementary Table S8, S8). The large number of differentially expressed genes in the parental generation comprises both sex-related genes and cultivar-specific differences. Genetic recombination in the F_2_ population randomized cultivar-specific traits unrelated to sex expression, allowing us to identify genes more directly associated with the monoecy trait. Consequently, to refine our transcriptomics analysis, we compared gene expression profiles between monoecious and dioecious female F_2_ plants, which reduced the number of differentially expressed genes to 90 (Fig. 4b, Supplementary Table S8, S9). However, none of these 90 genes were located within the *Monoecy1* QTL when applying the standard adjusted p-value filter (padj < 0.05) from DESeq2. This discrepancy likely reflects the limited sample size in our F_2_ transcriptomic analysis (n=3 per group), which resulted in high dispersion estimates and consequently many genes with NA adjusted p-values in DESeq2, despite some having extremely low unadjusted p-values.

**Figure 4.**
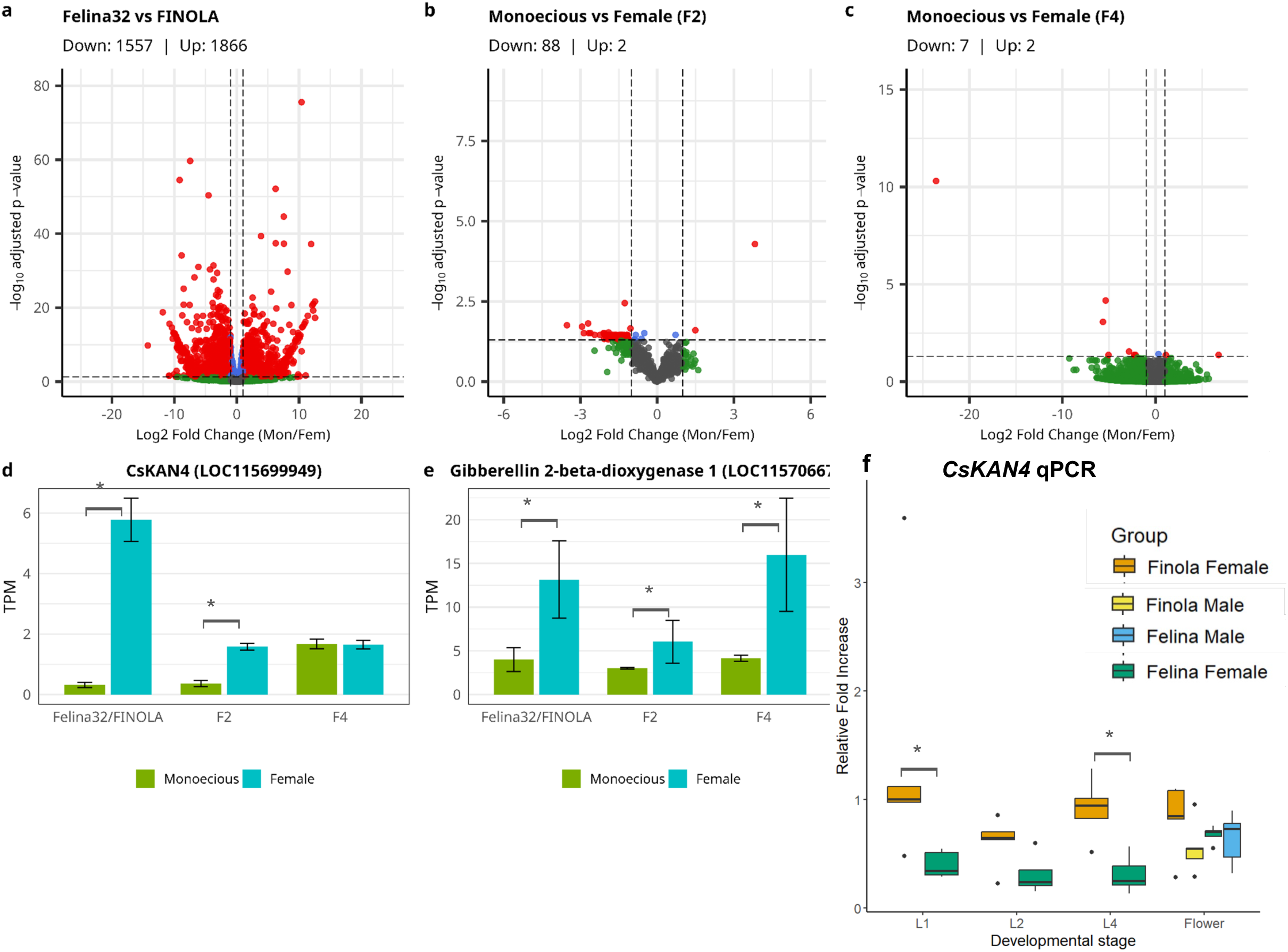
Transcriptomic analysis of monoecy across *Cannabis* generations. **a-c)** Volcano plots depicting differential gene expression between monoecious and female *C. sativa* plants across three generations. Log2 fold change (x-axis) represents expression in monoecious relative to female plants; -log10 adjusted p-value (y-axis) indicates statistical significance. Red dots represent significantly differentially expressed genes (DEGs; adjusted p < 0.05, |log2FC| > 1). **a)** Comparison between parental lines ‘Felina 32’ (monoecious) and ‘FINOLA’ (dioecious females) reveals extensive transcriptional differences. **b)** F_2_ segregating population shows a marked reduction in DEGs as genetic recombination has randomised cultivar-specific traits unrelated to sex expression. **c**) F_4_ generation demonstrates further reduction of DEGs. **d-f)** ETPM-normalized expression of key monoecy related candidate genes across generations and developmental stages. **d)** *Gibberellin 2-beta-dioxygenase 1* (LOC115706670), thought to be degrading GA, displays consistently downregulation in monoecious plants across generations. Error bars represent standard error of the mean. **e**) *CsKAN4* (LOC115699949) shows downregulation in leaf samples of monoecious parental line ‘Felina 32’ and in F_2_ monoecious plants compared to females, but not in flower meristem samples in F_4_ plants. **f)** *CsKAN4* (LOC115699949) qPCR showing is expressioned in the apical meristem in early stages of plant development (two leaf pairs, stage L2) and in later stages (four and nine leaf pairs, L4 and L9) and in flower tissue in cultivars ‘Felina 32’ (monoecious) and ‘FINOLA’ (dioecious). .* indicates p < 0.05. For panels d and e, p-values were obtained from DESeq2 pairwise group comparisons. For panel f, the Wilcoxon–Mann-Whitney test was used due to non-normality of the data.

To identify candidate genes within *Monoecy1* that might contribute to the monoecy trait, we employed a complementary approach: we examined genes that were both (1) significantly differentially expressed between the parental cultivars ‘FINOLA’ and ‘Felina 32’ (padj < 0.05, |log₂FC| > 1), and (2) showed consistent differential expression in the F_2_ generation using instead a p-value threshold (p < 0.01, |log₂FC| > 1). This intersection identified LOC115699949 as the only gene within *Monoecy1* meeting both criteria. LOC115699949 bears homologies to the *Arabidopsis thaliana* transcription factor gene *KANADI4* (*KAN4*/*ATS*) (Izhaki & Bowman, 2007; Gomez *et al*., 2016) and was therefore termed *CsKAN4* (Supplementary Fig. S13). *CsKAN4* is expressed more highly in female ‘FINOLA*’* than in monoecious ‘Felina 32’ plants, and also more highly in female F_2_ than in monoecious F_2_ plants (Fig. 3d). In the d_S_ analysis, *CsKAN4* was found to be present exclusively on the X chromosome with no corresponding homolog found on the Y chromosome.

We also extended our transcriptomic analysis to the F_4_ monoecious and dioecious samples of our mapping population, generated though random intercrossing of the F_2_ and F_3_ generation.

Here, the number of genes differentially expressed between monoecious and female dioecious plants was just 9 (Fig. 4c, Supplementary Table S8, S9).

We further identified nine genes consistently downregulated across all generations of our mapping population in monoecious vs. female dioecious plants when filtering for p < 0.05 and |log₂FC| > 1 (Supplementary Table S10), one of them being likely involved in the gibberellic acid (GA) pathway: *LOC115706670* (encoding a putative gibberellin 2-beta-dioxygenase 1), located on chromosome 1 (Fig. 4d). This enzyme is presumably involved in GA catabolism, suggesting that its reduced expression in monoecious plants may contribute to higher GA catabolism and consequently more elevated GA levels compared to dioecious females, thus promoting male flower development.

We did not detect differential expression of *CsKAN4* in the F_4_ generation (Fig. 4e), but F_4_ samples were isolated from flowering tissue whereas F_2_ and parental samples were from leaf tissue.To obtain a more detailed view of *CsKAN4*, we performed qPCR on apical meristem samples from monoecious and dioecious individuals at different vegetative stages and young flowering stages, using the *CsPP2A* housekeeping gene to normalize expression. This analysis confirmed significant downregulation of *CsKAN4* in monoecious ‘Felina 32’ plants compared to dioecious female ‘FINOLA’ plants (Fig. 4f, Supplementary Fig. S14), supporting the hypothesis that reduced *CsKAN4* expression is a molecular signature of the monoecious phenotype. Notably, *CsKAN4* differential expression was restricted to pre-flowering stages, consistent with the absence of differential expression observed in RNA-seq data from the inflorescences and flowers in F_4_ plants. Furthermore, *CsKAN4* expression did not differ significantly between male and female flowers of monoecious ‘Felina 32’ plants (Fig. 4f, (Supplementary Fig. S14).

### *Monoecy1* contains putative male vs. female sex-determination genes

Strikingly, a detailed analysis of the *CsKAN4* gene locus revealed that it is just 60,000 bp away from the ortholog of the *Arabidopsis* gene AT4G33280, *CsREM16*. In *Arabidopsis*, some genes of the REM (Reproductive Meristem) family are implicated in reproductive development (Mantegazza *et al*., 2014). For *Cannabis*, we previously reported that *CsREM16* was strongly upregulated in females as compared to males and proposed that it constitutes a molecular switch between male and female plant development (Shi *et al*., 2025). Like *CsKAN4*, *CsREM16* was found to have no corresponding Y chromosome counterpart.

We next analysed the region containing *CsREM16* and *CsKAN4* in more detail. Between *CsREM16* and *CsKAN4* only two other genes are located, a putative long noncoding (lnc) RNA (LOC133032448) and a putative pseudogene (LOC115699946) (Fig. 5a). Sequence analysis further revealed that the lncRNA contains a region of ∼300 nt with high similarity to *CsREM16* (Supplementary Fig. S15, Supplementary Data S2, Supplementary Text 2). This might indicate a possible regulatory function and we therefore termed LOC133032448 *lncREM16*.

**Figure 5.**
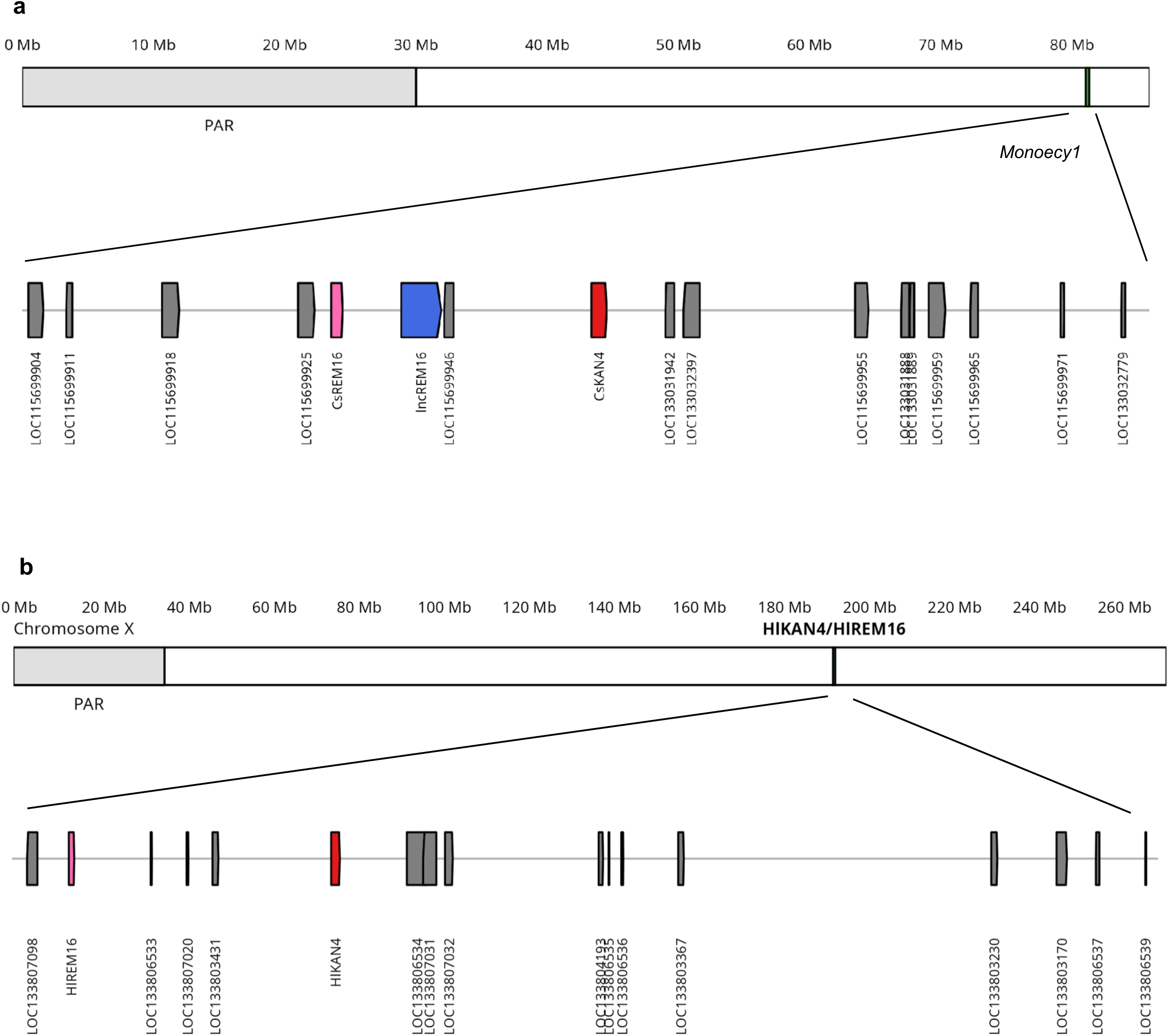
Genomic organization and expression patterns of the potential sex-determining genes in the *Monoecy1* locus. **a)** Chromosome X ideogram and detailed gene organization in the *Monoecy1* region surrounding *CsREM16* and *CsKAN4* (81.025-81.300 Mb). The upper panel shows the X chromosome with the pseudoautosomal region (PAR, gray) and the highlighted region containing potential sex-determining genes contained in the *Monoecy1* locus (green). The lower panel displays gene arrangements with directional arrows indicating strand orientation. The putative sex-determining genes are colored according to their sex-bias classification: *CsREM16* (pink, XX-biased), *lncREM16* (blue, male-biased), and *CsKAN4* (red, dioecious-biased). **b)** Genomic organization of candidate sex-determining gene orthologs on the *H. lupulus* X chromosome. *HlREM16* (LOC133803719) and *HlKAN4* (LOC133804030) are located in close proximity within the terminal non-recombining region of the X chromosome, as they are in *Cannabis*, at 189.9-190.0 Mb. Gene positions are indicated with directional arrows showing transcriptional orientation. The lncREM16 present in *Cannabis* was not identified in *H. lupulus* in its entirety.

To establish further evidence of the sex related expression pattern of the genes in *Monoecy1*, we integrated 64 RNA-seq samples from previous experiments, encompassing monoecious and dioecious samples from different developmental stages and growth conditions from the ‘FINOLA’ and’ ‘Felina 32’ cultivars (Dowling *et al*., 2024; Shi *et al*., 2025) (Supplementary Fig. S16a, Supplementary Table S11). This confirmed the dioecious-biased expression of *CsKAN4*, the high expression of *CsREM16* in female and monoecious individuals, and showed that *lncREM16* is largely silenced in monoecious and female dioecious individuals and remarkably, despite being located on the non recombining region of the X chromosome, *lncREM16* is expressed in male samples (Supplementary Fig. S16a, Supplementary Table S11). Notably, LOC115699965, which is among the genes with the highest d_S_ values and is homologous to the *Arabidopsis* pollen-function gene *JINGUBANG* (Ju *et al*., 2016) is situated approximately only 150kb downstream of our primary sex determination candidate *CsREM16* (Shi et al., 2025) and 90kb downstream of *CsKAN4*, further supporting the ancient origin of this locus. Furthermore, genome sequencing data from the monoecious ‘Felina 32’ cultivar revealed an insertion containing a transposable element domain ca. 8,000 bp upstream the start of the transcription site of *CsKAN4*, absent in the dioecious ‘FINOLA’ cultivar (Supplementary Fig. S17, Supplementary Data S3) as a possible disruptor of its expression in the monoecious cultivar.

### The candidate sex-determining genes are conserved between *Cannabis* and *Humulus*

Given that *Cannabis* and *Humulus* species separated after the evolution of their shared XY sex chromosomes and sex determination system in their ancestor (Prentout *et al*., 2021), we investigated whether the candidate sex-determining genes identified in *Cannabis* are conserved in hop (*Humulus lupulus*). Analysis of the hop X chromosome revealed that both *HlREM16* (LOC133803719) and *HlKAN4* (LOC133804030) are located in the terminal section of the non-recombining region (189.9-190 Mb), maintaining their close proximity as observed in *Cannabis* (Fig. 5b). Protein sequence comparison showed high conservation, with *HlREM16* sharing 83.3 % amino acid identity with *CsREM16*, while *HlKAN4* displayed a lower 59.3 % protein level identity with *CsKAN4* (Supplementary Fig. S18). The full *lncREM16* transcript was not identified in hop. Expression analysis using published RNA-seq data from five male and five female hop plants demonstrated that *HlREM16* exhibits the same female-biased expression pattern observed in hemp, with significantly higher (p < 0.05) expression in females. Similar to *CsKAN4*, *HlKAN4* showed a similar expression level (p > 0.05) in females compared to males (Supplementary Fig. S16b). In contrast to *Cannabis*, monoecy is uncommon in hops (Haunold, 1991) and thus a female vs monoecious *HlKAN4* expression comparison is not included here.

### Evaluation of alternative sex determination candidates

Given recent proposals of ethylene signaling components as sex determinants in *Cannabis* and *Humulus* (Akagi *et al*., 2025; Carey *et al*., 2025), we evaluated the expression patterns of the two key proposed sex determination candidates: the ethylene biosynthesis gene *Cannabis sativa 1-aminocyclopropane-1-carboxylic acid synthase 3* (*CsACS3*, *LOC115696400*) (Carey *et al*., 2025) and the putative ethylene receptor gene *Cannabis sativa Ethylene Receptor 1* (*CsETR1*, *LOC115702329*) (Akagi *et al*., 2025). Both genes reside within the terminal non-recombining region of the X chromosome, positioning them as plausible sex determination factors according to our model. Both are, however, not in the *Monoecy1* QTL, though *CsACS3* is located very close to the QTL, at 79.96 Mb, while the QTL is positioned from 80.8Mb to 82.1 Mb.

To enable direct comparison of gene expression dynamics, we examined our three primary candidates (*CsREM16*, *CsKAN4*, and *lncREM16*) across the same developmental stages and tissues as the ethylene pathway genes (*CsACS3* and *CsETR1*). In vegetative apical meristem samples as well as in flowers *CsREM16* showed consistent female-biased expression (Fig. 6a). *lncREM16* displayed the opposite pattern, with male-specific expression in vegetative and flowering samples (Fig. 6b). *CsKAN4* showed similar expression in both sexes during vegetative and flowering stages with no significant male-female difference, consistent with its proposed role in the monoecy/dioecy distinction rather than male/female determination (Fig. 6c).

**Figure 6.**
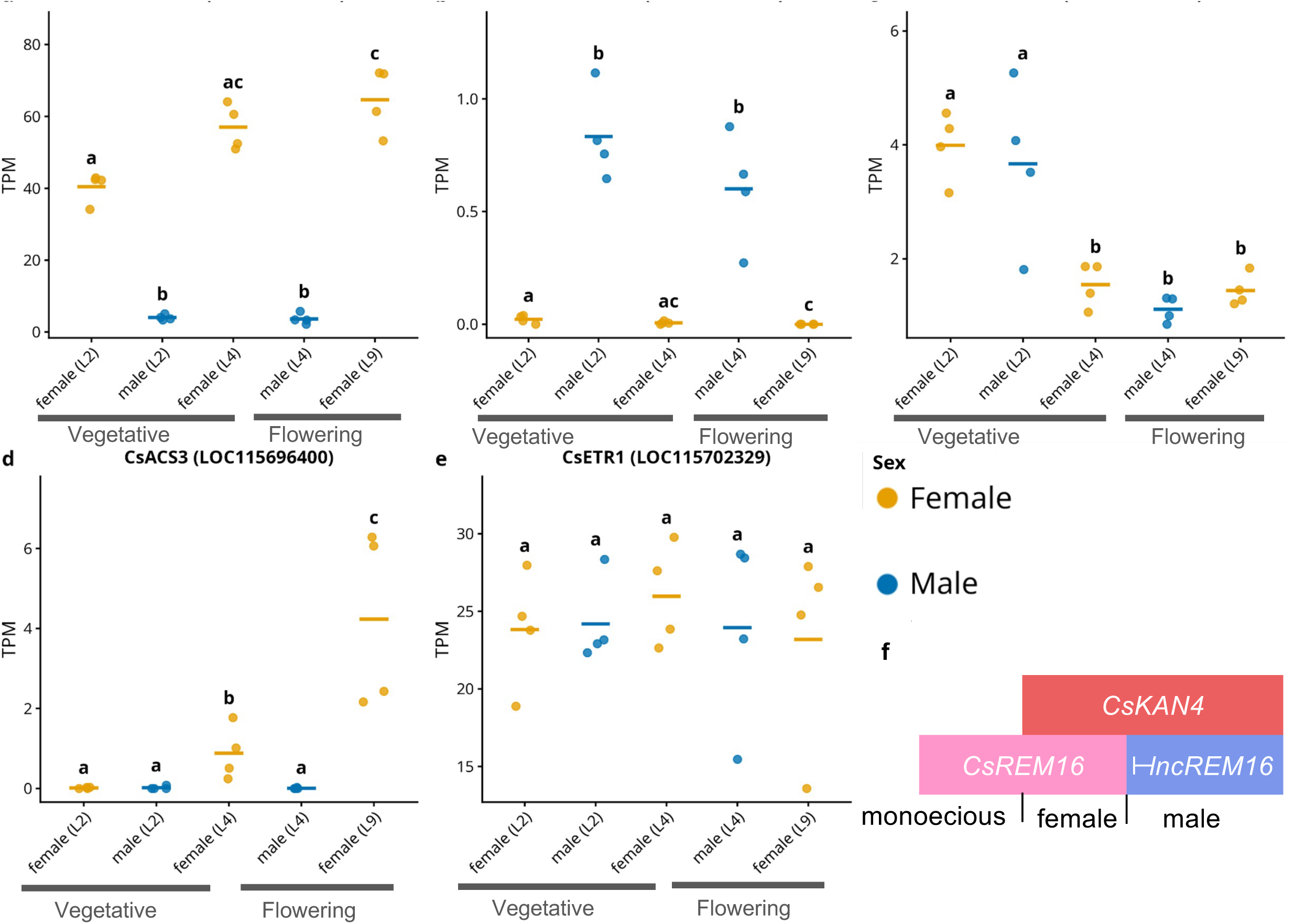
Expression analysis of sex determination candidate genes across developmental stages and cultivars. **(a–c)** Expression of primary sex-determination candidates in apical meristem samples across developmental stages. **(a)** *CsREM16* (LOC115699937) shows consistent female-biased expression at all stages examined **(b)** lncREM16 (LOC133032448) displays male-specific expression with no detectable expression (TPM = 0) in females. **(c)** CsKAN4 (LOC115699949) shows similar expression between sexes across all developmental stages. **(d–e)** Expression of alternative proposed ethylene signaling pathway sex determinants in apical meristem samples. **(d)** CsACS3 (LOC115696400) shows no or very low expression at early vegetative stages (L2) and in flowering males (L4), with detectable expression emerging in late vegetative females (L4) and flowering females (L9), albeit at low levels (TPM ∼1–4). **(e)** CsETR1 (LOC115702329) exhibits constitutive expression across all developmental stages and sex phenotypes, with no significant differences between males and females (adjusted p-value > 0.05). Expression values are shown as transcripts per million (TPM). Statistical significance was assessed using DESeq2 pairwise contrasts with Benjamini-Hochberg adjusted p-values. Groups sharing the same letter are not significantly different (adjusted p < 0.05). Orange: female; blue: male. Horizontal bars indicate group means. Developmental stages are two (L2), four (L4) and 9 (L9) leaf pair stages. Shoot apical meristem or flowering tissues were sampled. Males flower at L4, females at L9. Stages and RNAseq data from Shi *et al*. (2025) **f)** Combinatorial model of sex determination in Cannabis governed by three key regulators: CsREM16, CsKAN4 and lncREM16. XX females express CsREM16 and CsKAN4 but not lncREM16, resulting in female flower development. XY males express lncREM16 and CsKAN4 but CsREM16 is silenced (potentially by lncREM16), leading to male flower development. XX monoecious plants express CsREM16 but not lncREM16 and CsKAN4, enabling development of both male and female flowers on the same individual.

Analysis of *CsACS3* expression across our dataset revealed tissue and developmental specificity. The gene shows no or very low expression at the vegetative stage and in flowering males, but expression is detectable in late-stage vegetative female and flowering female samples, albeit at low levels (TPM ∼ 1 to 4) (Fig. 6d). *CsACS3* has also been proposed as a candidate gene for controlling monoecy vs. dioecy, with two indels in exon 4 being associated with monoecy (Carey et al., 2025). Sequence analysis of the *CsACS3* locus revealed no polymorphisms between the monoecious ‘Felina 32’ and dioecious ‘FINOLA’ plants used to generate our mapping population across the entire protein coding regions (Supplementary Fig. S19).

The proposed sex determining ethylene receptor gene *CsETR1* displayed a different expression dynamic. Unlike the tissue- and sex-specific *CsACS3*, *CsETR1* maintained similar expression across all developmental stages and sex phenotypes examined (Fig. 6e, Supplementary Fig. S16), with mean TPM values ranging from 18 to 27.

Similarly, comparison between dioecious female and males ‘FINOLA’ and monoecious ‘Felina 32’ across a variety of tissues and sampling conditions revealed no significant expression differences for either *CsACS3* and *CsETR1* (adjusted p-value > 0.05) (Supplementary Fig. S16c,d).

We next reasoned that a master switch for sex determination would be expected not to change in expression even if the phenotypic sex is altered by phytohormone/hormone inhibitor application. To further analyse this we leveraged a transcriptomic data set by (Adal *et al*., 2021) that provides expression data from relatively mature male (from XY plants) and female (XX plants) flowers but also male flowers that were induced by silver thiosulfate on female plants (i.e. male flowers on XX plants). Expression of *CsREM16, CsACS3* and *lncREM16* is associated with the genotypic sets in those datasets (Supplementary Fig. S20), i.e. *CsREM16* and *CsACS3* are expressed in female (XX) and induced male (XX) flowers at higher levels than in male (XY) flowers (for *CsETR1* the pattern was similar though not statistically significant, potentially due to high dispersion). For *lncREM16* expression in induced male flowers (XX) remains zero, similar to female flowers (XX), whereas some expression is detected in male flowers (XY). *CsKAN4* showed low expression in all three groups, indicating again low expression in the late flowering stage (Supplementary Fig. S20).

## Discussion

Taken together, our data indicate that one of the oldest parts of the non-recombining region of the *Cannabis* X chromosome contains the *Monoecy1* QTL which contributes to the development of monoecious vs. dioecious individuals. *Monoecy1* contains three genes in the space of just 60,000 bp, the combinatorial expression of which predicts the sex of *Cannabis* plants: in monoecious individuals *CsREM16* is expressed, in female individuals *CsREM16* and *CsKAN4* are expressed and in male individuals *lncREM16* and *CsKAN4* are expressed (Fig. 6f).

We propose that *CsREM16, CsKAN4* and *lncREM16* constitute molecular determinants of both monoecy/dioecy and male/female sex expression in *Cannabis*. CsREM16 and CsKAN4 are transcription factors belonging to the B3 and GARP family respectively, and homologs of those proteins in *Arabidopsis* have been demonstrated to be involved in reproductive development (Mantegazza *et al*., 2014; Safi *et al*., 2017). Previous studies have indicated that sex determination in *Cannabis* is under X chromosome gene dosage control (Akagi *et al*., 2025), which is consistent with the X chromosomal location and male or female biased expression of the genes identified here. *CsREM16* and *lncREM16* are not affected in their expression by the masculinizing chemical silver thiosulfate, consistent with the idea that they are very high in the hierarchy of the sex determination cascade, possibly upstream of phytohormone pathway. Furthermore, that several genes controlling sex-determination are found on the oldest evolutionary stratum on the *Cannabis* X chromosome supports the canonical model for the evolution of plant sex chromosomes (Charlesworth, 2013), as also found recently in *Silene latifolia* (Moraga *et al*., 2025).

Though the exact molecular mechanisms of how the three genes control sex determination remains elusive, it is interesting to note that *kan4* mutants in *Arabidopsis* show an increased expression of GA biosynthesis genes and an increased level of GA (Gomez *et al*., 2016). Likewise, our transcriptomic data indicate that in monoecious individuals a gene encoding a putative gibberellin 2-beta-dioxygenase 1 (*GA2ox1*) is downregulated in conjunction with *CsKAN4*. GA2ox1 enzymes are involved in GA catabolism. Intriguingly, exogenous GA applications induce the formation of male flowers on female plants in *Cannabis* (Ram & Jaiswal, 1972) and a gene involved in the GA pathway has previously been implicated in sex determination in *Cannabis* (Petit *et al*., 2020). A reduction of *CsKAN4* expression in monoecious plants may thus lead to an increase in GA, which in turn may foster the development of male flowers. We detected differential *CsKAN4* expression mainly in vegetative tissue, including leaves. Long distance transport of GA has been demonstrated in other plants (Eriksson *et al*., 2006; Regnault *et al*., 2015b,a). It thus appears conceivable that transcriptional changes in *CsKAN4* in vegetative organs can exert effects on floral development.

The molecular role of *CsREM16* in sex determination is more difficult to pinpoint, but our data indicate that its expression is strongly associated with XX as opposed to XY individuals across different developmental stages and cultivars (Fig. 4b) (Shi *et al*., 2025). In *Glycine max* the *REM16* ortholog *GmREM16* is involved in controlling flowering time (Singh Yadav, 2025; Wang *et al*., 2025). In *Arabidopsis*, some *REM* mutants display strong female gametophytic defects (Matias-Hernandez *et al*., 2010; Mendes *et al*., 2016; Caselli *et al*., 2019). Furthermore, recent data from *Arabidopsis* indicate that REM transcription factors recruit RNA polymerase IV to activate expression of small interfering RNAs that in turn initiate DNA methylation of specific loci (Wu *et al*., 2025; Xu *et al*., 2025). Taken together, this may suggest that *CsREM16* has been co-opted to control sex determination of the sporophyte via changing the DNA methylation landscape.

In contrast to other plant sex determination genes, which are predominantly expressed in reproductive tissues (Leite Montalvão *et al*., 2021; Masuda & Akagi, 2023; Charlesworth & Harkess, 2024; Marais *et al*., 2025), *CsREM16* is expressed in all analysed female tissues. This is arguably unusual for a plant sex determination gene but a similar system exists e.g. in *Drosophila* where every somatic cell in females expresses the sex determination gene *SEX LETHAL* which is not expressed in males (Salz & Erickson, 2010). In *Drosophila*, sex is specified in a cell autonomous manner, and it will be fascinating to see whether a similar concept may apply to *Cannabis*.

Interestingly, *HlREM16*, the ortholog of *CsREM16* in *Humulus* (hops), is also located on the X chromosome and upregulated in female vs. male individuals. *Humulus* is the closest extant relative of *Cannabis* and also dioecious, and the sex chromosomes in the two genera originated before the separation of the lineages leading to hop and *Cannabis* (Prentout *et al*., 2021). It is thus conceivable that the two genera share a similar sex determination system and that *HlREM16* and *CsREM16* are key regulators of female identity in both species. The conservation of gene location, sequence, and expression patterns between *Cannabis* and *Humulus* provides evidence that this sex-determining system predates their divergence, supporting the ancient origin of these sex chromosomes and the fundamental role of these genes in Cannabaceae sex determination.

Interestingly, in *Cannabis* we detected with *lncREM16* a long non-coding RNA with an expression pattern opposite to that of *CsREM16* and with sequence similarities to *CsREM16.* We consider it plausible that there is a regulatory relationship between the two genes, in which one may repress the other or they repress each other in a feedback loop.

We were not able to detect the long noncoding RNA *lncREM16* in hop, which may be taken as indication that the details of the sex determination system in hop and *Cannabis* diverged over time. Similar observations have been made in poplar, where two closely related species share the same gene regulatory circuits governing sex determination but the molecular details of gene regulation differ (Müller *et al*., 2020).

### Quo vadis, sex determination?

A number of different (not necessarily mutually exclusive) models have been proposed recently on how sex expression and sex determination in the Cannabaceae is controlled (Shi *et al*., 2025; Akagi *et al*., 2025; Carey *et al*., 2025; Monthony *et al*., 2026). Each of them makes testable predictions about the role of different X and Y chromosomal genes.

Two of the proposed sex determination genes are involved in ethylene biosynthesis or perception: *CsETR1* and *CsACS3* (Akagi *et al*., 2025; Carey *et al*., 2025). While ethylene clearly plays an important role in *Cannabis* sex expression, with male plants producing female flowers after ethephon treatment and female plants producing male flowers after silver nitrate (an ethylene inhibitor) treatment (Ram & Jaiswal, 1972; Mohan Ram & Sett, 1982; Monthony *et al*., 2026), it is noteworthy that ethylene seems to change the sex expression of the flowers but not the overall difference in architecture of male vs female plants (Shi *et al*., 2024; Garcia-de Heer *et al*., 2025; Monthony *et al*., 2026). Furthermore, whereas *CsACS3* is upregulated in female plants starting at the four leaf pair stage, broad transcriptomic differences between male and female plants are already detected earlier, with tens of genes overexpressed in females, and hundreds overexpressed in males already at the 2-leaf pair stage (Shi *et al*., 2025). *CsETR1* did show differential expression between male and female plants only in mature flowers at relatively late developmental stages, but not earlier during development (Figure 6, Supplementary Figure S20). Taken together, we hypothesize that *CsETR1* and *CsACS3* might well be involved in controlling sex expression and determining male vs. female floral identity. However, they may not be the master switches controlling sex determination. The phenotypic difference between male and female *Cannabis* plants extends beyond floral morphology, and we predict sex determination genes like the putative candidate *CsREM16* act upstream of *CsETR1* and *CsACS3*. Functional analyses will likely be required to elucidate the regulatory relationships between those genes and to understand whether they act independently from each other or in one regulatory cascade.

A polymorphism, specifically deletions in the fourth exon of *CsACS3,* was also proposed as a candidate for the distinction between monoecy and dioecy. However, the amino acid sequence of *CsACS3* in the monoecious ‘Felina 32’ and dioecious ‘FINOLA’ plants used in our mapping population are identical, indicating that, though CsACS3 might be involved in sex determination or sex expression, an amino acid sequence polymorphism in this gene may not be the sole causal mutation for the monoecy/dioecy trait in ‘FINOLA’ vs ‘Felina 32’. More broadly, the discrepancies between our and previous findings (Carey *et al*., 2025) may reflect distinct genetic origins of monoecy across cultivars.

Moreover, the incomplete penetrance observed in our segregation analysis supports a polygenic model of monoecy/dioecy sex determination. *Monoecy1* explains approximately 15 % of phenotypic variance for monoecy/dioecy, suggesting additional genetic factors influence sex expression. These may include multiple loci selected through breeding, environmental interactions, and epigenetic regulation. There is a high degree of phenotypic variation in sex expression even within individual monoecious cultivars (Faux *et al*., 2014; Trubanová *et al*., 2025), supporting the notion that monoecy is under intricate genetic and environmental control. Further exploration of the link between GA homeostasis and sex expression may shed additional light on this complex phenotype and its genetic control.

In conclusion, all of the sex determination candidates discussed here (*CsREM16*, *lncREM16*, *CsKAN4*, *CsACS3* and *CsETR1*) may therefore be involved in controlling sex expression and sex determination to some extent. It will be fascinating to see how future experiments shape our understanding of sex determination in *Cannabis* and beyond.

## Acknowledgements

This publication has emanated from research conducted with the financial support of Taighde Éireann – Research Ireland under grant number IRCLA/2022/3294. CAD was supported by an Irish Research Council–Environmental Protection Agency Government of Ireland Postgraduate Scholarship (grant no. GOIPG/2019/1987).

## Author contributions

MT conducted the study performing the bioinformatics analyses and generated figures and text. AM and ARB conducted sampling, DNA, and RNA extractions and molecular analyses. CD designed and created the original mapping population used as the basis for the experiment. SS and RM supervised the work. QR analyzed the strata with help from RCRLV. TG provided input in the interpretations. RM and MT wrote the manuscript with input from SS. RCRLV QR and TG commented on the manuscript. All authors read and approved the manuscript.

## Data availability

All relevant data are contained in the manuscript and supplementary files.

## Supplementary Material

### Supplementary Tables

**Table S1. d_s_ and d_n_ values for the X-Y single copy orthologs in the Kompolti’ cultivar**

**Table S2. Sex phenotype segregation across three generations of *Cannabis sativa* crosses and maternal inheritance patterns**

**Table S3. Genetic linkage map and quantitative trait locus (QTL) analysis for sex determination using standard interval mapping**

**Table S4. Refined QTL mapping for sex determination using composite interval mapping analysis**

**Table S5. Association between marker X_80375543 genotypes and sex phenotype distribution in F_2_ female and monoecious plants**

**Table S6. Validation of male sex phenotype through Y chromosome read mapping analysis**

**Table S7. Gene content and sex-biased expression patterns in the oldest evolutionary stratum of the X chromosome (80-86 Mb)**

**Table S8. Differential gene expression analysis between monoecious and dioecious *Cannabis* plants across parental and segregating generations**

**Table S9. Significantly differentially expressed genes (adjusted p-value < 0.05, |log2FC| > 1) between monoecious and dioecious plants across generations. For the F2 a p-value <0.01 filter was used instead of the adjusted p-value due to the amount of NAs values.**

**Table S10. Core set of genes consistently downregulated in monoecious plants across multiple generations**

**Table S11. Expression validation of sex determination candidate genes across 64 RNA-seq samples from parental cultivars (BioProject PRJNA956491)**

### Supplementary Figures

**Figure S1. Changepoint analysis for inferring evolutionary strata based on synonymous divergence (d_S_) between X and Y chromosomes in *Cannabis* and *Humulus*.**

**Figure S2. Progeny sex distribution by maternal phenotype.**

**Figure S3. F_2_ population genotype matrix.**

**Figure S4. Genetic map and marker linkage analysis.**

**Figure S5. Genotype distribution by sex phenotype at X chromosome markers.**

**Figure S6. Genetic map after removal of unexpected genotypes.**

**Figure S7. QTL analysis with filtered genotype data.**

**Figure S8. QTL analysis after filtering the mapped reads for MAPQ >35.**

**Figure S9. QTL analysis on data mapped and genotyped against ‘Santhica’ haplotypes.**

**Figure S10. Genotypes and phenotypes at QTL region of interest.**

**Figure S11. Gene loss analysis using alternative reference genome.**

**Figure S12. Y-specific gene content distribution.**

**Figure S13. Phylogenetic reconstruction of KAN4/ATS homologs.**

**Figure S14. CsKAN4 qPCR using alternative housekeeping gene**

**Figure S15. Sequence similarity between *lncREM16* and *CsREM16***

**Figure S16. Gene expression of the candidate sex-determining genes**

**Figure S17. Transposable element presence in monoecious cultivars ‘Santhica 27’ and ‘Felina 32’.**

**Figure S18. Protein sequence conservation between *CsREM16* (LOC115699937) / *HlREM16* (LOC133803719) and between *CsKAN4* / *HlKAN4* (LOC133804030)**

**Figure S19. Sequence conservation of the ethylene biosynthesis gene *CsACS3* (LOC115696400) between monoecious and dioecious cultivars.**

**Figure S20 Expression of the sex determination candidate gene in male, female, and silver-induced male flowers.**

### Supplementary Data

**Supplementary Data S1** Single tree phylogenies of the 313 genes where single-copy orthologs between *Cannabis* X and Y and *Humulus* X and Y chromosomes were identified. Lists of the trees with the two possible topologies, distinguishing whether recombination, were identified are included.

**Supplementary Data S2** Nucleotide alignment of *lncREM16* showing sequence similarity to *CsREM16*.

**Supplementary Data S3** Nucleotide sequence of the transposable element found upstream of the *CsKAN4* gene in the monoecious ‘Felina 32’ parent but not the dioecious ‘FINOLA’ parent.

### Supplementary Text

**Supplementary Text 1** Analysis of unexpected X chromosome genotypes in the F_2_ mapping population.

**Supplementary Text 2** Characterization of the male-specific long non-coding RNA *lncREM16* and its sequence relationship to *CsREM16*.

## References

Adal AM, Doshi K, Holbrook L, Mahmoud SS. 2021. Comparative RNA-Seq analysis reveals genes associated with masculinization in female Cannabis sativa. Planta 253: 1–17.

Adhikary D, Kulkarni M, El-Mezawy A, Mobini S, Elhiti M, Gjuric R, Ray A, Polowick P, Slaski JJ, Jones MP, et al. 2021. Medical Cannabis and Industrial Hemp Tissue Culture: Present Status and Future Potential. Frontiers in Plant Science 12: 627240.

Akagi T, Segawa T, Uchida R, Tanaka H, Shirasawa K, Yamagishi N, Yaegashi H, Natsume S, Takagi H, Abe A, et al. 2025. Evolution and functioning of an X–A balance sex-determining system in hops. Nature Plants: 1–14.

Andre CM, Hausman J-F, Guerriero G. 2016. Cannabis sativa: The Plant of the Thousand and One Molecules. Frontiers in Plant Science 7.

Bachtrog D. 2013. Y chromosome evolution: emerging insights into processes of Y chromosome degeneration. Nature reviews. Genetics 14: 113.

Bergero R, Charlesworth D. 2009. The evolution of restricted recombination in sex chromosomes. Trends in Ecology & Evolution 24: 94–102.

Broman KW, Wu H, Sen Ś, Churchill GA. 2003. R/qtl: QTL mapping in experimental crosses. Bioinformatics 19: 889–890.

Carey SB, Bentz PC, Lovell JT, Akozbek LM, Myers ZA, Korani W, Havill JS, Padgitt-Cobb L, Lynch RC, Allsing N, et al. 2025. An X-linked sex determination mechanism in cannabis and hop. bioRxiv: 2024.12.09.627636.

Caselli F, Beretta VM, Mantegazza O, Petrella R, Leo G, Guazzotti A, Herrera-Ubaldo H, de Folter S, Mendes MA, Kater MM, et al. 2019. REM34 and REM35 Control Female and Male Gametophyte Development in Arabidopsis thaliana. Frontiers in Plant Science 10.

Charlesworth D. 2013. Plant sex chromosome evolution. Journal of Experimental Botany 64: 405–420.

Charlesworth D, Harkess A. 2024. Why should we study plant sex chromosomes? The Plant Cell 36: 1242–1256.

Danecek P, Auton A, Abecasis G, Albers CA, Banks E, DePristo MA, Handsaker RE, Lunter G, Marth GT, Sherry ST, et al. 2011. The variant call format and VCFtools. Bioinformatics 27: 2156–2158.

DePristo MA, Banks E, Poplin R, Garimella KV, Maguire JR, Hartl C, Philippakis AA, del Angel G, Rivas MA, Hanna M, et al. 2011. A framework for variation discovery and genotyping using next-generation DNA sequencing data. Nature Genetics 43: 491–498.

Divashuk MG, Alexandrov OS, Razumova OV, Kirov IV, Karlov GI. 2014. Molecular Cytogenetic Characterization of the Dioecious Cannabis sativa with an XY Chromosome Sex Determination System. PLOS ONE 9: e85118.

Dowling CA, Shi J, Toth JA, Quade MA, Smart LB, McCabe PF, Schilling S, Melzer R. 2024. A FLOWERING LOCUS T ortholog is associated with photoperiod-insensitive flowering in hemp (Cannabis sativa L.). The Plant Journal 119: 383–403.

Emms DM, Kelly S. 2019. OrthoFinder: phylogenetic orthology inference for comparative genomics. Genome Biology 20: 238.

Ewels P, Magnusson M, Lundin S, Käller M. 2016. MultiQC: summarize analysis results for multiple tools and samples in a single report. Bioinformatics 32: 3047–3048.

Faux A-M, Berhin A, Dauguet N, Bertin P. 2014. Sex chromosomes and quantitative sex expression in monoecious hemp (Cannabis sativa L.). Euphytica 196: 183–197.

Garcia-de Heer L, Guo Q, Mieog J, Nolan M, Liu L, Dimopoulos N, Melzer R, Kretzschmar T. 2025. Transcriptomic analysis of ethephon-induced sex reversion of male Cannabis sativa reveals changes in expression of floral homeotic genes and a distinct trichome morphology. Journal of Experimental Botany 76: 5401–5418.

Garcia-de Heer L, Mieog J, Burn A, Kretzschmar T. 2024. Why not XY? Male monoecious sexual phenotypes challenge the female monoecious paradigm in Cannabis sativa L. Frontiers in Plant Science 15: 1412079.

Gomez MD, Ventimilla D, Sacristan R, Perez-Amador MA. 2016. Gibberellins Regulate Ovule Integument Development by Interfering with the Transcription Factor ATS. Plant Physiology 172: 2403.

Guo R, Guo H, Zhang Q, Guo M, Xu Y, Zeng M, Lv P, Chen X, Yang M. 2018. Evaluation of reference genes for RT-qPCR analysis in wild and cultivated Cannabis. Bioscience, Biotechnology, and Biochemistry 82: 1902–1910.

Harkess A, Zhou J, Xu C, Bowers JE, Van Der Hulst R, Ayyampalayam S, Mercati F, Riccardi P, McKain MR, Kakrana A, et al. 2017. The asparagus genome sheds light on the origin and evolution of a young Y chromosome. Nature Communications 8: 1279.

Haunold A. 1991. 29 - Cytology and Cytogenetics of Hops. Developments in Plant Genetics and Breeding Volume 2, Part B: 551–563.

Hyden B, Zou J, Wilkerson DG, Carlson CH, Robles AR, DiFazio SP, Smart LB. 2023. Structural variation of a sex-linked region confers monoecy and implicates GATA15 as a master regulator of sex in Salix purpurea. New Phytologist 238: 2512–2523.

Izhaki A, Bowman JL. 2007. KANADI and Class III HD-Zip Gene Families Regulate Embryo Patterning and Modulate Auxin Flow during Embryogenesis in Arabidopsis. The Plant Cell 19: 495–508.

Ju Y, Guo L, Cai Q, Ma F, Zhu Q-Y, Zhang Q, Sodmergen. 2016. Arabidopsis JINGUBANG Is a Negative Regulator of Pollen Germination That Prevents Pollination in Moist Environments. The Plant Cell 28: 2131–2146.

Leite Montalvão AP, Kersten B, Fladung M, Müller NA. 2021. The Diversity and Dynamics of Sex Determination in Dioecious Plants. Frontiers in Plant Science 11.

Lindeløv J. 2020. mcp: An R Package for Regression With Multiple Change Points.

Love MI, Huber W, Anders S. 2014. Moderated estimation of fold change and dispersion for RNA-seq data with DESeq2. Genome Biology 15: 550.

Lubell JD, Brand MH. 2018. Foliar Sprays of Silver Thiosulfate Produce Male Flowers on Female Hemp Plants. HortTechnology 28: 743–747.

Lynch RC, Padgitt-Cobb LK, Garfinkel AR, Knaus BJ, Hartwick NT, Allsing N, Aylward A, Bentz PC, Carey SB, Mamerto A, et al. 2025. Domesticated cannabinoid synthases amid a wild mosaic cannabis pangenome. Nature 643: 1001–1010.

Mantegazza O, Gregis V, Mendes MA, Morandini P, Alves-Ferreira M, Patreze CM, Nardeli SM, Kater MM, Colombo L. 2014. Analysis of the arabidopsis REM gene family predicts functions during flower development. Annals of Botany 114: 1507–1515.

Marais GAB, Branco C, Rocheta M, Dufay M, Tonnabel J. 2025. Plant sex-determining genes and the genetics of the evolution towards dioecy. Journal of Experimental Botany 76: 3896–3911.

Margarido GRA, Souza AP, Garcia A a. F. 2007. OneMap: software for genetic mapping in outcrossing species. Hereditas 144: 78–79.

Masuda K, Akagi T. 2023. Evolution of sex in crops: recurrent scrap and rebuild. Breeding Science 73: 95–107.

Matias-Hernandez L, Battaglia R, Galbiati F, Rubes M, Eichenberger C, Grossniklaus U, Kater MM, Colombo L. 2010. VERDANDI is a direct target of the MADS domain ovule identity complex and affects embryo sac differentiation in Arabidopsis. The Plant Cell 22: 1702–1715.

Mendes MA, Guerra RF, Castelnovo B, Silva-Velazquez Y, Morandini P, Manrique S, Baumann N, Groß-Hardt R, Dickinson H, Colombo L. 2016. Live and let die: a REM complex promotes fertilization through synergid cell death in Arabidopsis. Development 143: 2780–2790.

Mohan Ram HY, Sett R. 1982. Induction of fertile male flowers in genetically female Cannabis sativa plants by silver nitrate and silver thiosulphate anionic complex. Theoretical and Applied Genetics 62: 369–375.

Monthony AS, Roy J, Ronne M de, Carlson O, Murch SJ, Torkamaneh D. 2026. Sex-specific ethylene responses drive floral sexual plasticity in Cannabis sativa. The Plant Journal 125: e70721.

Moraga C, Branco C, Rougemont Q, Jedlička P, Mendoza-Galindo E, Veltsos P, Hanique M, Vega RCR de la, Tannier E, Liu X, et al. 2025. The Silene latifolia genome and its giant Y chromosome. Science (New York, N.Y.) 387: 630.

Müller NA, Kersten B, Leite Montalvão AP, Mähler N, Bernhardsson C, Bräutigam K, Carracedo Lorenzo Z, Hoenicka H, Kumar V, Mader M, et al. 2020. A single gene underlies the dynamic evolution of poplar sex determination. Nature Plants 6: 630–637.

Pancaldi F, Salentijn EMJ, Trindade LM. 2025. From fibers to flowering to metabolites: unlocking hemp (Cannabis sativa) potential with the guidance of novel discoveries and tools. Journal of Experimental Botany 76: 109–123.

Petit J, Salentijn EMJ, Paulo M-J, Denneboom C, Trindade LM. 2020. Genetic Architecture of Flowering Time and Sex Determination in Hemp (Cannabis sativa L.): A Genome-Wide Association Study. Frontiers in Plant Science 11: 569958.

Prentout D, Stajner N, Cerenak A, Tricou T, Brochier-Armanet C, Jakse J, Käfer J, Marais GAB. 2021. Plant genera Cannabis and Humulus share the same pair of well-differentiated sex chromosomes. New Phytologist 231: 1599–1611.

Ram HYM, Jaiswal VS. 1972. Induction of male flowers on female plants of Cannabis sativa by gibberellins and its inhibition by abscisic acid. Planta 105: 263–266.

Riera-Begue A, Toscani M, Malik A, Dowling C, Schilling S, Melzer R. 2025. A simple and reliable PCR-based method to differentiate between XX and XY sex genotypes in Cannabis sativa. bioRxiv: 2025.05.02.651914.

Rougemont Q, Lucotte E, Boyer L, Jalaber de Dinechin A, Snirc A, Giraud T, Rodríguez de la Vega RC. 2025. EASYstrata: an all-in-one workflow for genome annotation and genomic divergence analysis. NAR Genomics and Bioinformatics 7.

Ryu B-R, Gim G-J, Shin Y-R, Kang M-J, Kim M-J, Kwon T-H, Lim Y-S, Park S-H, Lim J-D. 2024. Chromosome-level Haploid Assembly of Cannabis sativa L. cv. Pink Pepper. Scientific Data 11: 1442.

Safi A, Medici A, Szponarski W, Ruffel S, Lacombe B, Krouk G. 2017. The world according to GARP transcription factors. Current Opinion in Plant Biology 39: 159–167.

Salz H, Erickson JW. 2010. Sex determination in Drosophila: The view from the top. Fly 4: 60–70.

Schilling S, Dowling CA, Shi J, Ryan L, Hunt DJL, O’Reilly E, Perry AS, Kinnane O, McCabe PF, Melzer R. 2021. The Cream of the Crop: Biology, Breeding, and Applications of Cannabis Sativa. : 471–528.

Shi J, Schilling S, Melzer R. 2024. Morphological and genetic analysis of inflorescence and flower development in hemp (Cannabis sativa L.). : 2024.01.25.577276.

Shi J, Toscani M, Dowling CA, Schilling S, Melzer R. 2025. Identification of genes associated with sex expression and sex determination in hemp (Cannabis sativa L.). Journal of Experimental Botany 76: 175–190.

Singh Yadav A. 2025. Time to bloom: GmREM16a promotes flowering time in soybeans. Plant Physiology 198.

Timoteo Junior AA, Oswald IWH. 2024. Optimized guidelines for feminized seed production in high-THC Cannabis cultivars. Frontiers in Plant Science 15: 1384286.

Trubanová N, Isobe S, Shirasawa K, Watanabe A, Kelesidis G, Melzer R, Schilling S. 2025. Genome-specific association study (GSAS) for exploration of variability in hemp (Cannabis sativa). Scientific Reports 15: 8371.

Vasimuddin Md, Misra S, Li H, Aluru S. 2019. Efficient Architecture-Aware Acceleration of BWA-MEM for Multicore Systems. In: 2019 IEEE International Parallel and Distributed Processing Symposium (IPDPS). Rio de Janeiro, Brazil: IEEE, 314–324.

Wang Z, Du C, Li Q, Li M, Wang Y, Bao G, Yin Y, Yang M, Yang Q, Xu P, et al. 2025. The transcription factor REM16a promotes flowering time in soybean by activating flowering-related genes. Plant Physiology 198.

Wu Z, Xue Y, Wang S, Shih Y-H, Zhong Z, Feng S, Draper J, Lu A, Hoeke CA, Sha J, et al. 2025. REM transcription factors and GDE1 shape the DNA methylation landscape through the recruitment of RNA polymerase IV transcription complexes. Nature Cell Biology 27: 1136–1147.

Xu G, Chen Y, Martins LM, Li E, Wang F, Magana T, Ruan J, Law JA. 2025. Transcription factors instruct DNA methylation patterns in plant reproductive tissues. Nature Cell Biology 27: 2116–2127.

Yamano K, Haseda A, Iwabuchi K, Osabe T, Sudo Y, Pachakkil B, Tanaka K, Suzuki Y, Toyoda A, Hirakawa H, et al. 2024. QTL analysis of femaleness in monoecious spinach and fine mapping of a major QTL using an updated version of chromosome-scale pseudomolecules. PLOS ONE 19: e0296675.

Yang Z. 2007. PAML 4: Phylogenetic Analysis by Maximum Likelihood. Molecular Biology and Evolution 24: 1586–1591.

Zhao Y, Liu Z, She H, Xu Z, Zhang H, Zheng S, Qian W. 2025. Comparative Transcriptome Analysis of Gene Expression Between Female and Monoecious Spinacia oleracea L. Genes 16: 24.

