## Supplementary Figures for "An ancient X chromosomal region harbours three genes potentially controlling sex determination in *Cannabis sativa*"

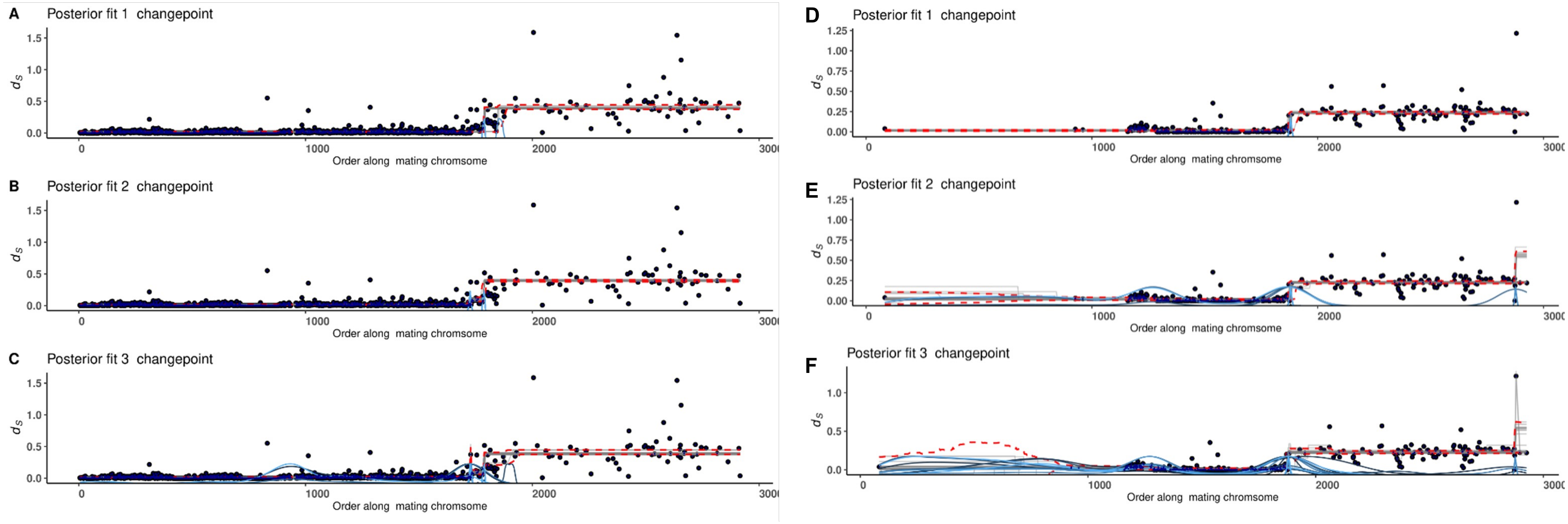


**Figure S1. Changepoint analysis for inferring evolutionary strata based on synonymous divergence ($d_S$) between X and Y chromosomes in *Cannabis* and *Humulus*.** Per-gene $d_S$ values for single-copy orthologs plotted along the X chromosome gene order, with Bayesian changepoint analysis used to infer evolutionary strata. **(A-C)** *Cannabis sativa* models with 1, 2, and 3 changepoints. **(D-F)** *Humulus lupulus* models with 1, 2, and 3 changepoints. Blue curves at the bottom of the x-axis represent posterior distributions of changepoint locations; dashed red lines indicate 2.5% and 97.5% quantiles of fitted values. Model comparison using leave-one-out cross-validation supported the single changepoint in both species. The similar location of the primary changepoint in both species suggests that recombination suppression across most of the non-recombining region predates the *Cannabis*-*Humulus* divergence.

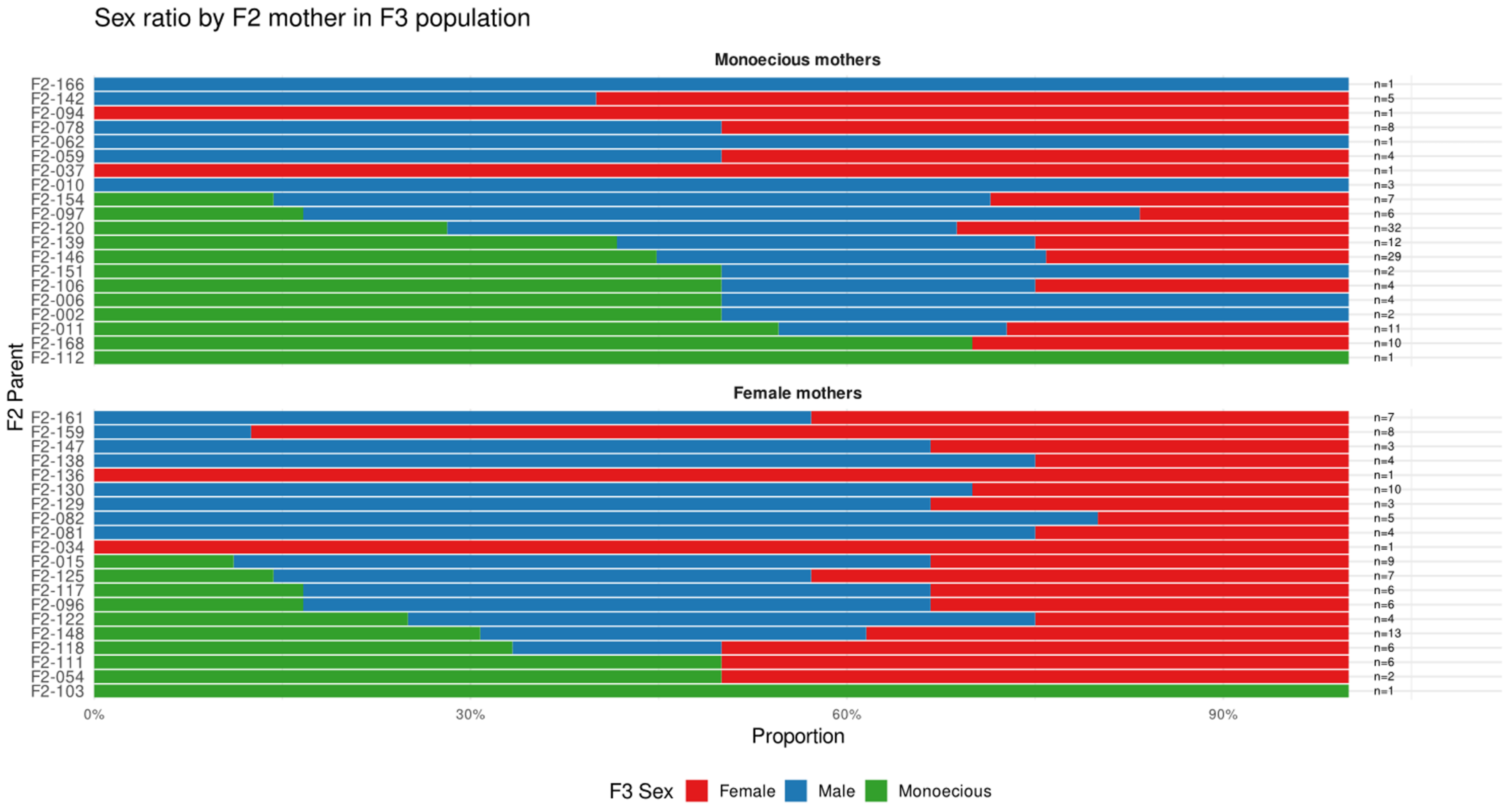


**Figure S2. Progeny sex distribution by maternal phenotype.**

Sex ratios of F3 progeny grouped by F2 maternal genotype (monoecious or female). Each bar represents offspring from a single F2 mother plant (identified on y-axis). Random pollination occurred within the F2 population. Monoecious F2 mothers consistently produced higher proportions of monoecious offspring compared to female F2 mothers, which predominantly yielded female progeny.

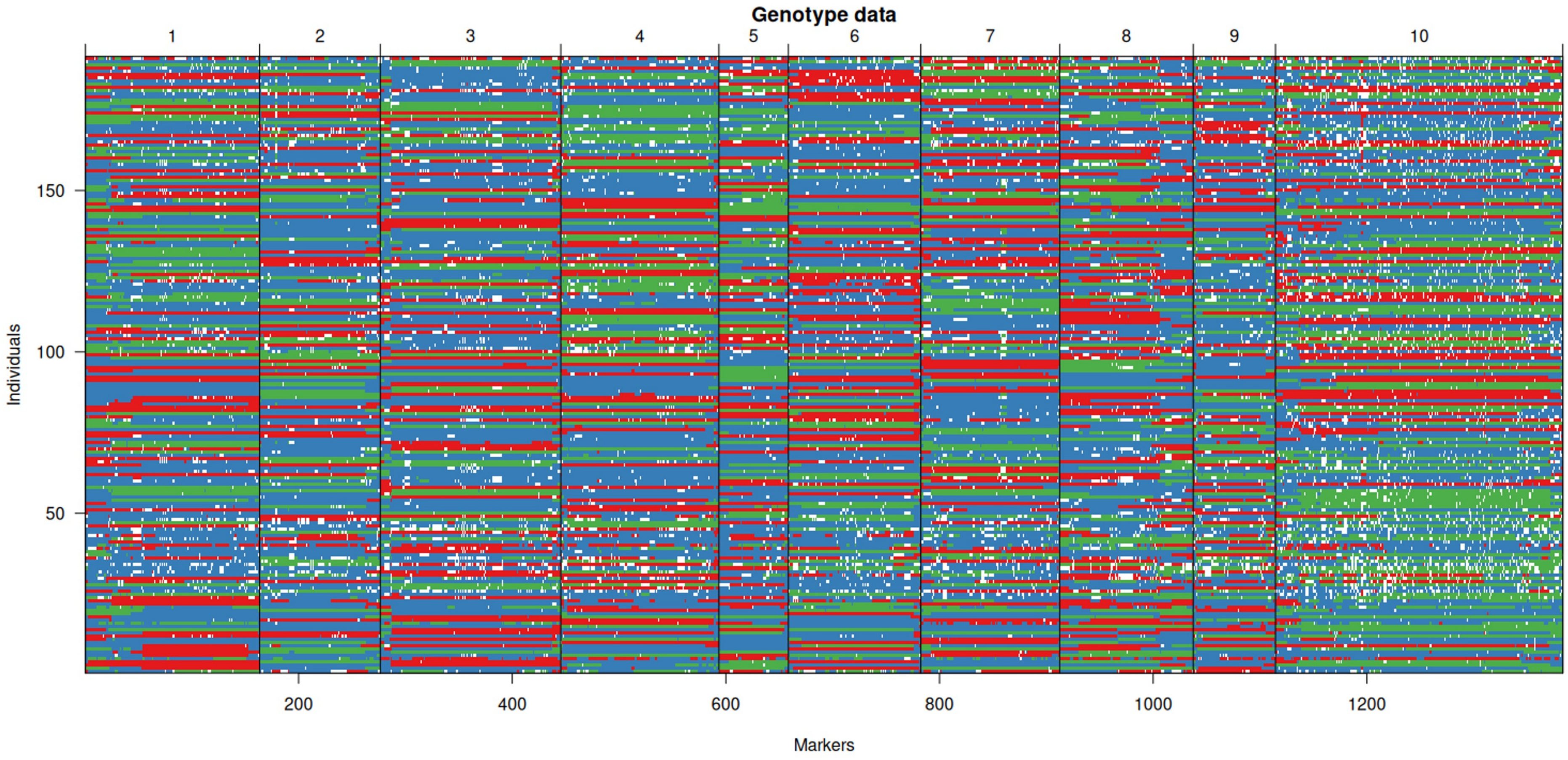


**Figure S3. $F_2$ population genotype matrix.**

Visualization of genotypes across all $F_2$ individuals (rows) and markers (columns) after marker reordering based on recombination frequencies. Color coding: red (homozygous 'FINOLA'), blue (heterozygous), green (homozygous 'Felina 32'), and white (missing data). Marker clustering within chromosomes demonstrates genetic linkage.

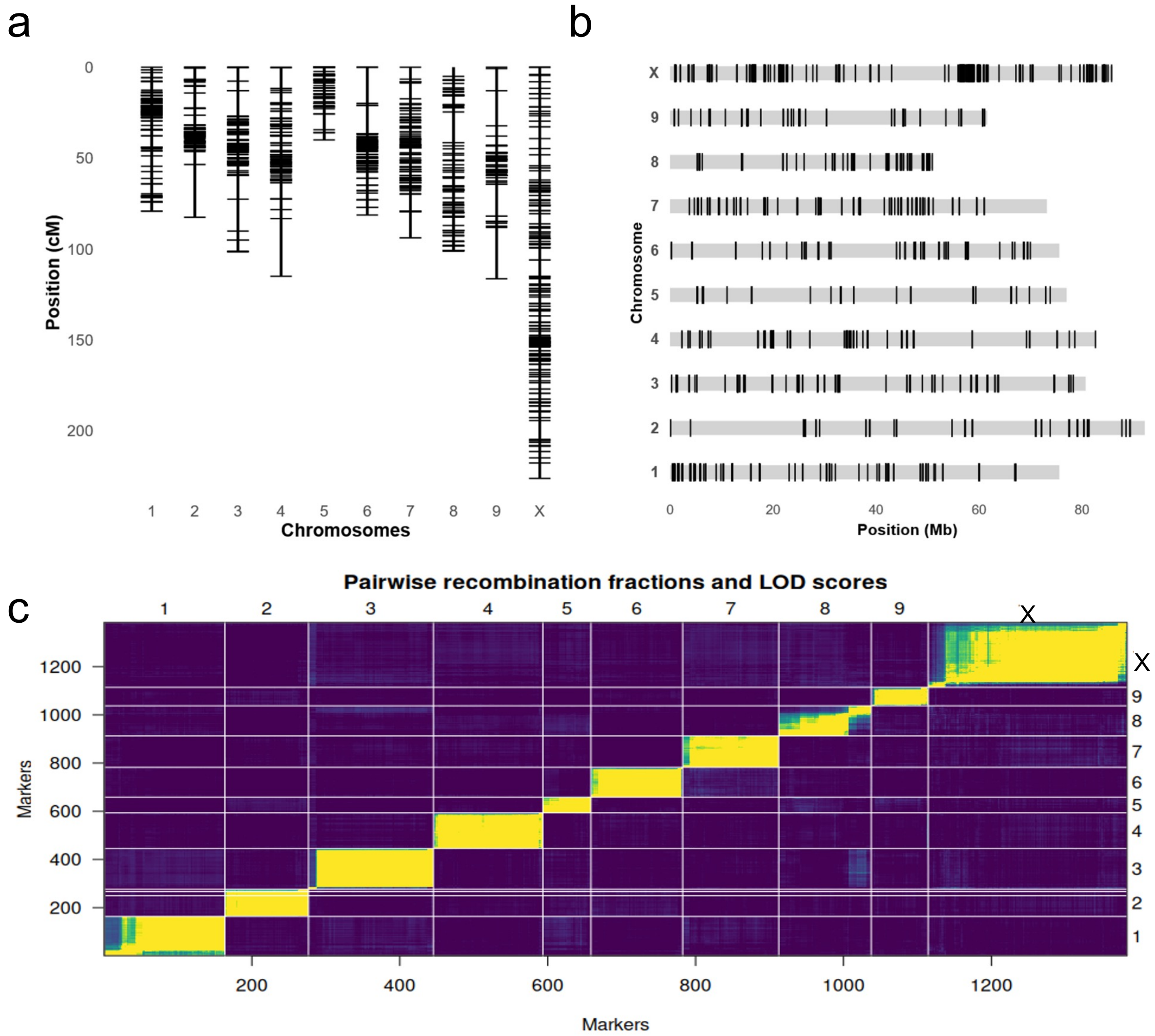


**Figure S4. Genetic map and marker linkage analysis.** a) Genetic map constructed from 1,383 high-quality SLAF-seq markers spanning 1,055 centiMorgans (cM) across 10 chromosomes, with an average marker spacing of < 1 cM. Chromosomes 1-9 represent autosomes, and chromosome X represents the sex chromosome. b) Physical map showing marker distribution along the chromosomes in megabases (Mb). The X chromosome spans approximately 86 Mb on the reference genome.
c)Pairwise marker relationships displayed as a heat map. Upper-left triangle: recombination fractions between markers (brighter shades indicate lower recombination frequencies, suggesting physical proximity). Lower-right triangle: LOD scores for marker linkage (brighter colors indicate stronger statistical evidence for linkage). The distinct diagonal blocks confirm accurate marker assignment to chromosomes.

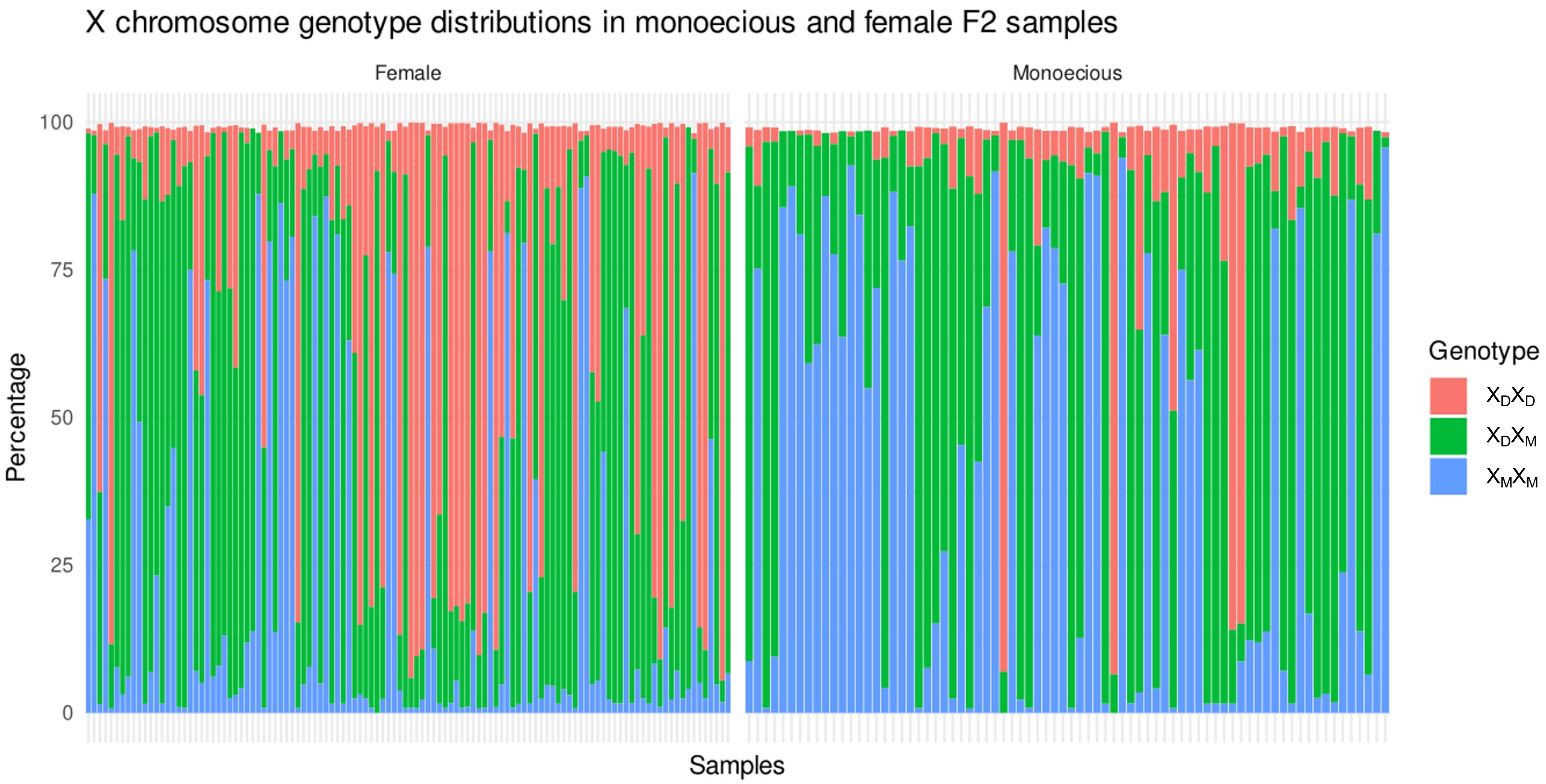


**Figure S5. Genotype distribution by sex phenotype at X chromosome markers.**
Frequency distribution of genotypes ($X_DX_D$: homozygous FINOLA, $X_DX_M$: heterozygous, $X_MX_M$: homozygous Felina 32) across X chromosome markers in F2 individuals, separated by sex phenotype. Female plants show enrichment for $X_DX_D$ genotypes while monoecious plants display higher prevalence of $X_MX_M$ genotypes, supporting X-linked inheritance of the monoecy trait.

a

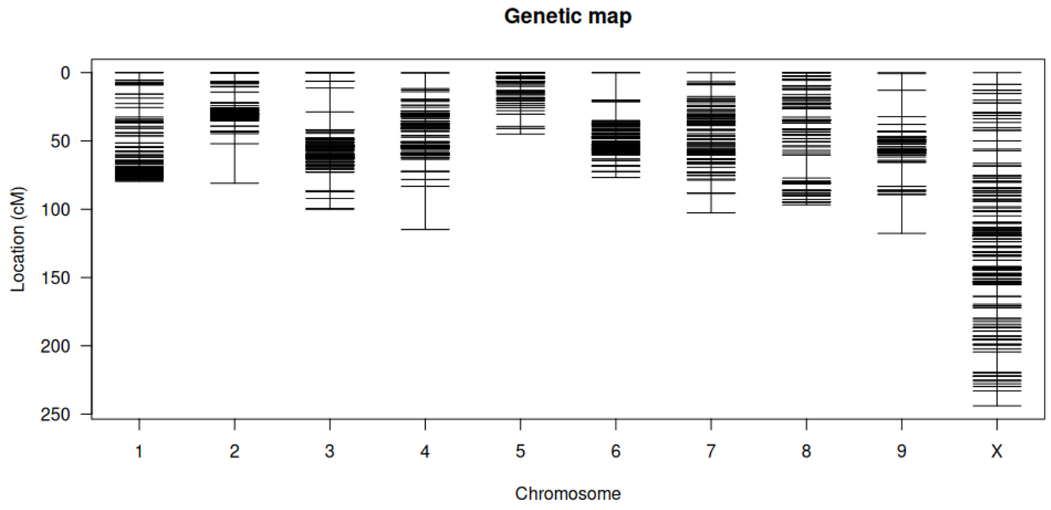


b

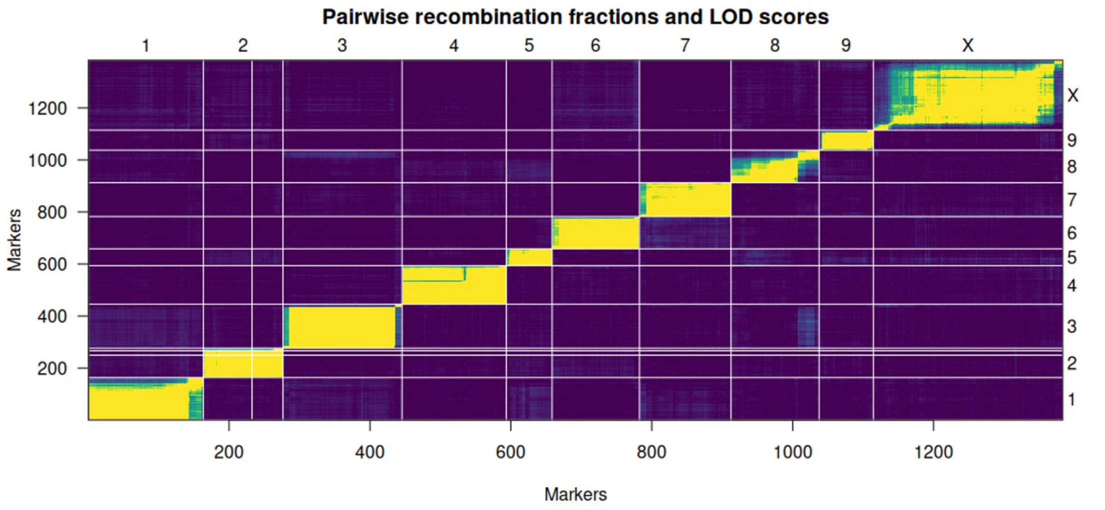


**Figure S6. Genetic map after removal of unexpected genotypes.**

a) Genetic map and linkage analysis after setting unexpected X chromosome genotypes (homozygous 'FINOLA' in female plants) to missing values. Marker order and genetic distances remain largely unchanged compared to the complete dataset, demonstrating robustness of the genetic map. b) Recombination fractions and LOD scores show genetic map coherence after removing the $X_DX_D$ genotypes from the X chromosome.

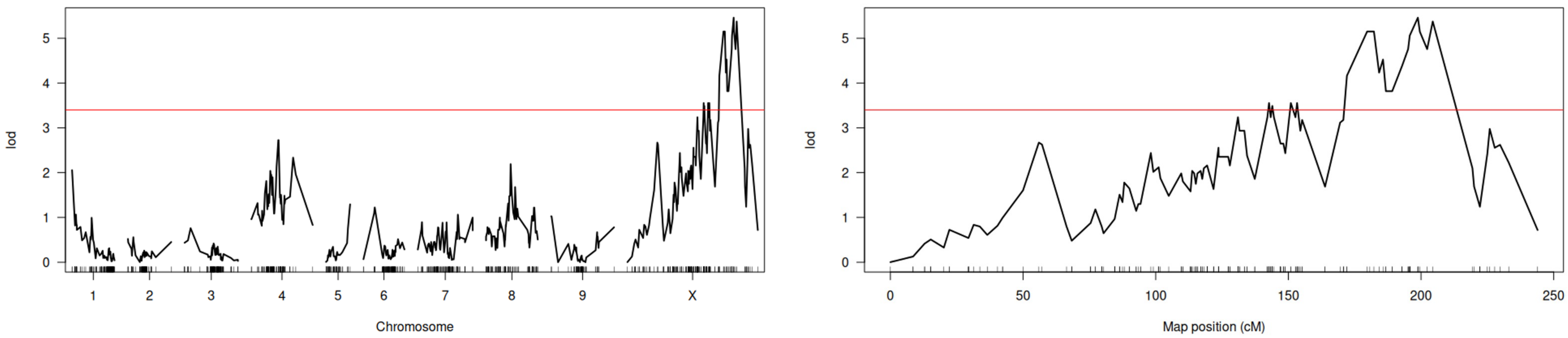


**Figure S7. QTL analysis with filtered genotype data.**
Monoecy trait QTL scan after removal of homozygous 'FINOLA' $X_DX_D$ chromosome genotypes. While overall LOD significance is reduced, the peak location remains consistent with the original analysis, confirming the robustness of the *Monoecy1* locus identification.

**Figure S8. Genotypes and phenotypes at QTL region of interest.**

Genotypes in red ($X_DX_D$, homozygous 'FINOLA'), blue ($X_DX_M$, heterozygous), green ($X_MX_M$, homozygous 'Felina 32'), and white (missing data) plotted for markers in the QTL region of interest for F2 monoecious and female plants, and corresponding plant phenotypes.

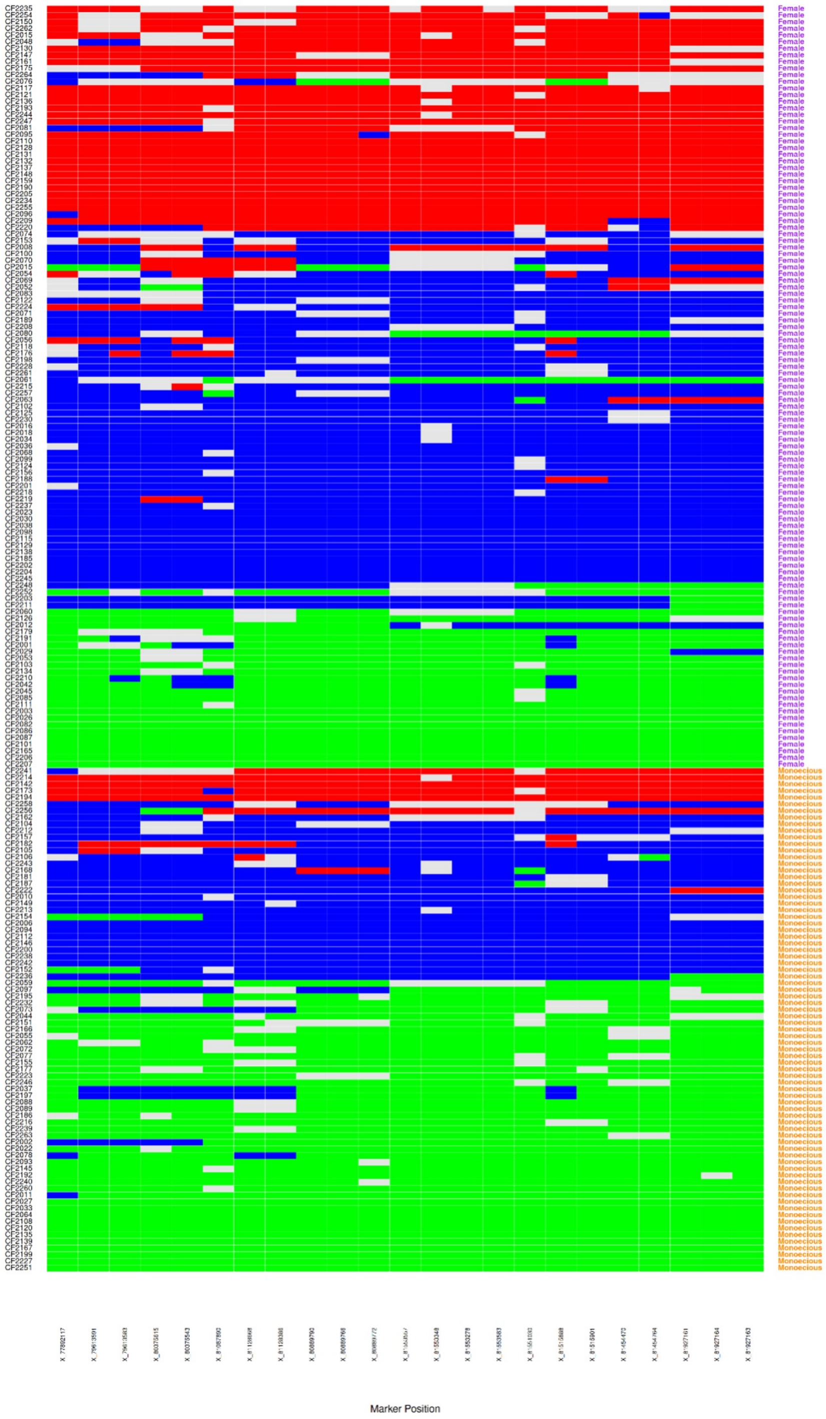

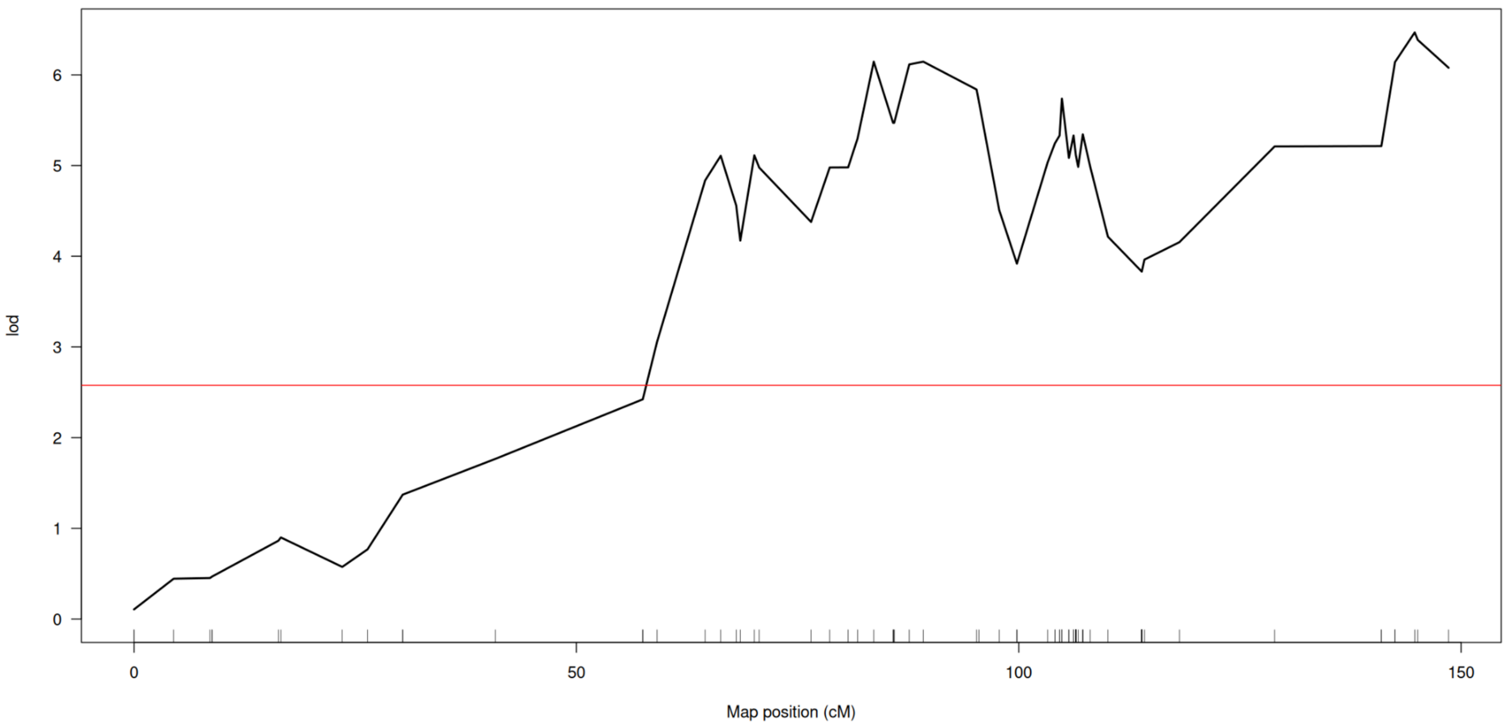


**Figure S9. QTL analysis after filtering the mapped reads for MAPQ >35.**
Monoecy trait QTL scan after removal of reads with mapping quality <35 . While overall marker number is reduced, the peak location remains consistent with the original analysis, confirming the robustness of the *Monoecy1* locus identification.

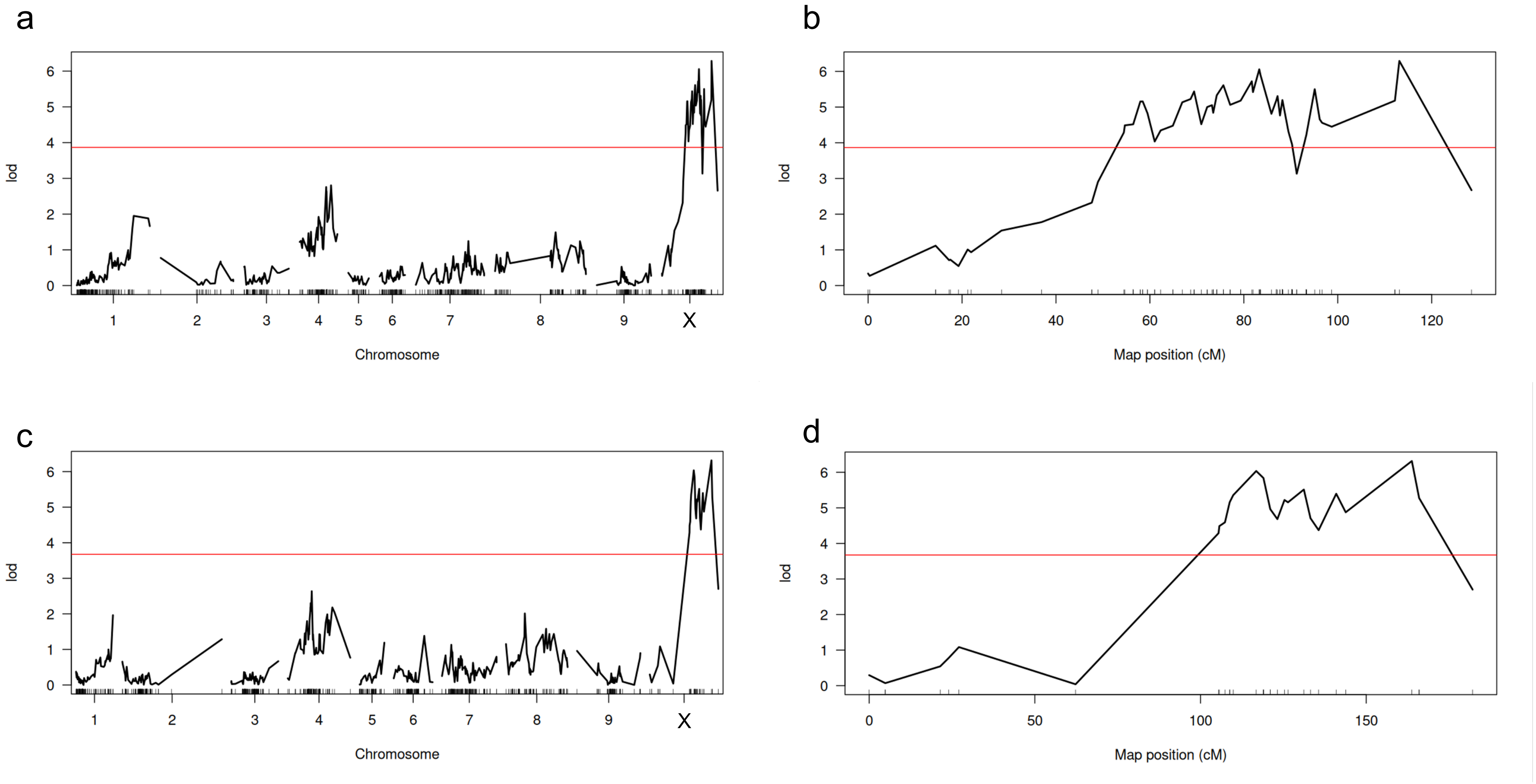


**Figure S10. QTL analysis on data mapped and genotyped against Santhica haplotypes.**
Monoecy trait QTL scan after mapping the SLAF-seq reads against the SAN2a (a, b) and SAN2b (c-d) monoecious Santhica assemblies. The peak location remains consistent with the original analysis against the Pink Pepper reference genome when mapping against both Santhica haplotypes, confirming the robustness of the *Monoecy1* locus identification.

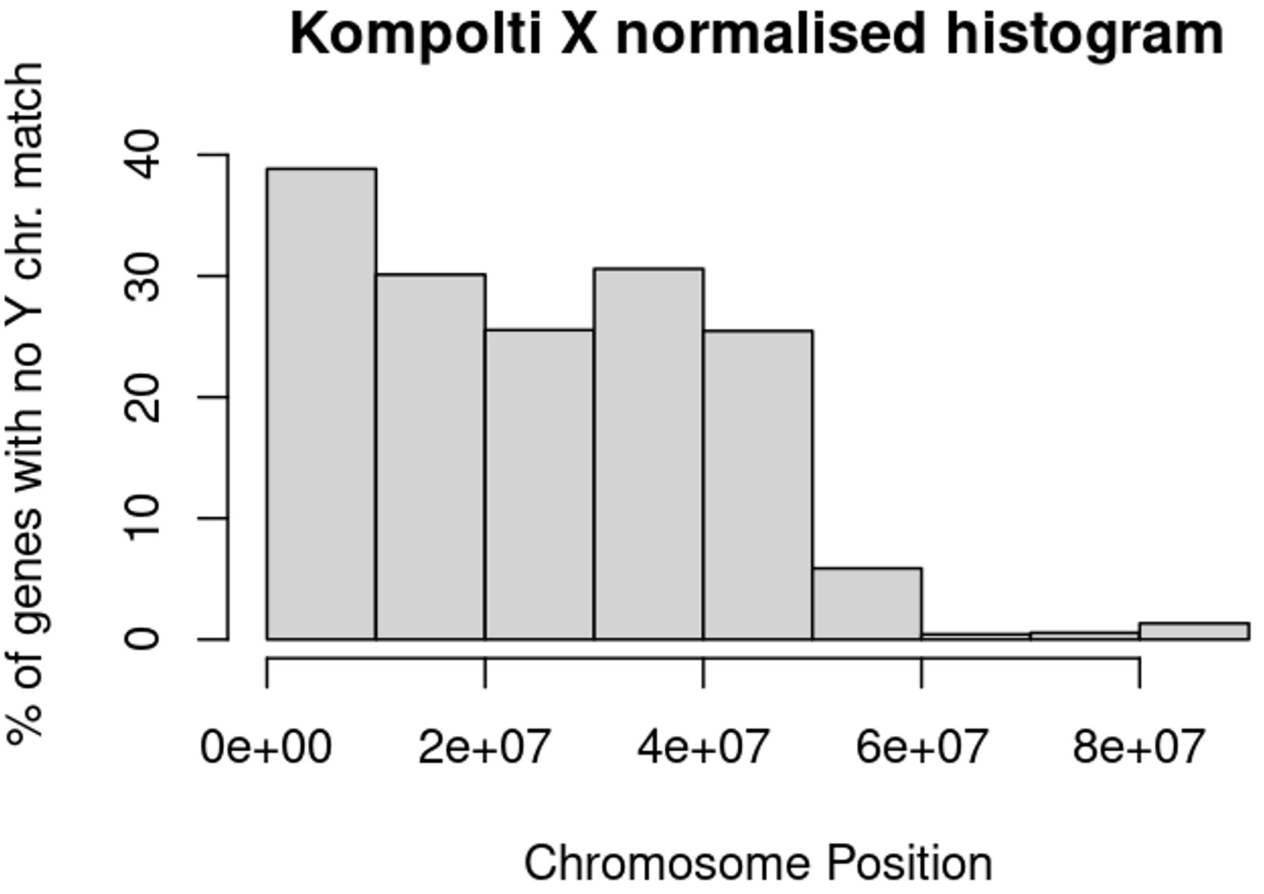


**Figure S11. Gene loss analysis using alternative reference genome.**
Percentage of X-chromosome hemizygous genes (genes without Y chromosome homologs) across X chromosome segments using the Kompolti cultivar reference genome. The pattern is similar to findings from the Pink Pepper reference, with increasing divergence from the pseudoautosomal region toward the chromosome terminus. The sequencing order of the Kompolti X chromosome is reversed when compared to Pink Pepper, the pseudoautosomal region is towards the end of the chromosome here.

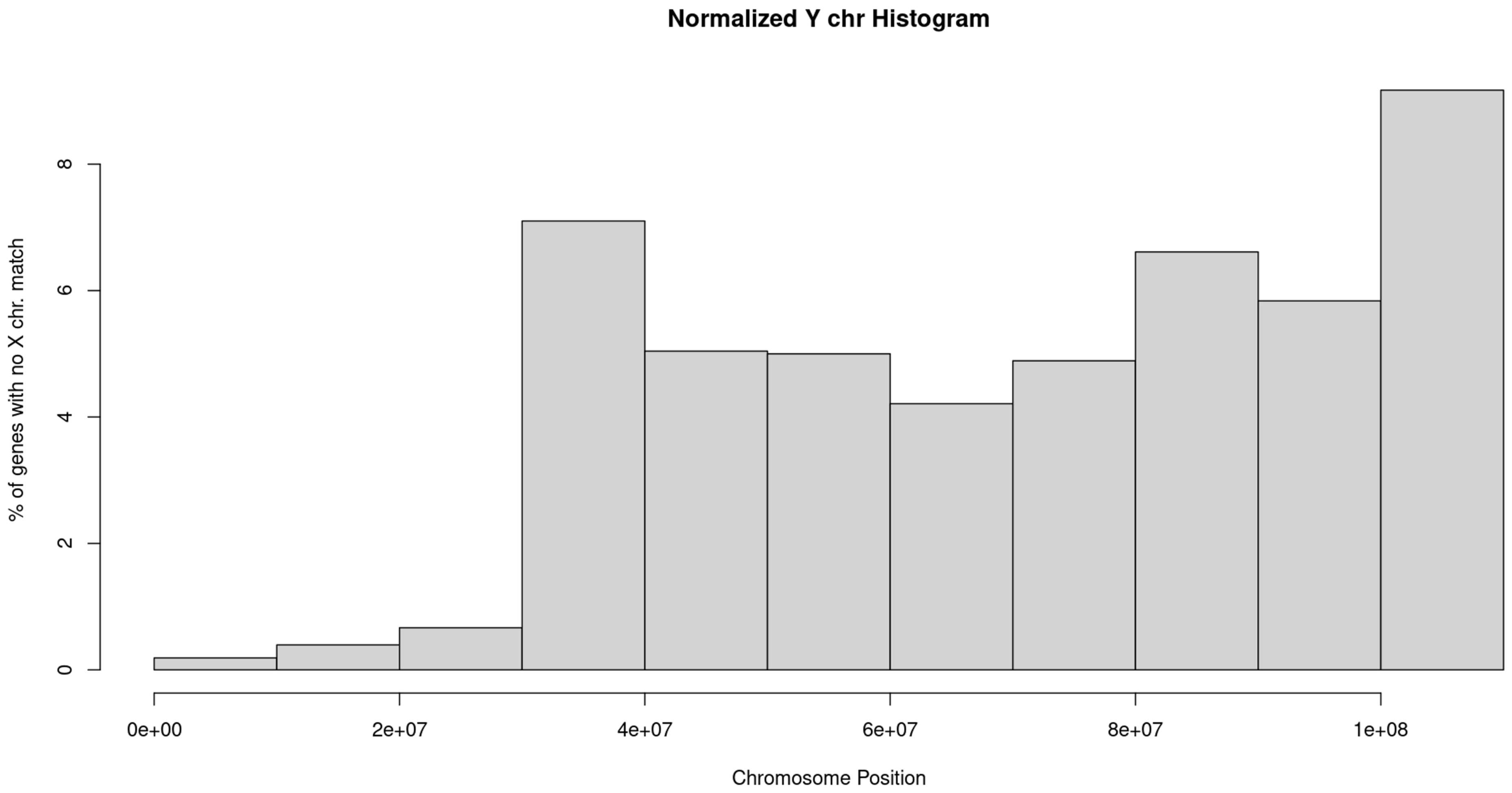


**Figure S12. Y-specific gene content distribution.**
Percentage of Y-chromosome hemizygous genes (genes without X chromosome homologs) across Y chromosome segments. Lower percentages compared to X-hemizygous genes reflect the reduced gene content of the degenerating Y chromosome.

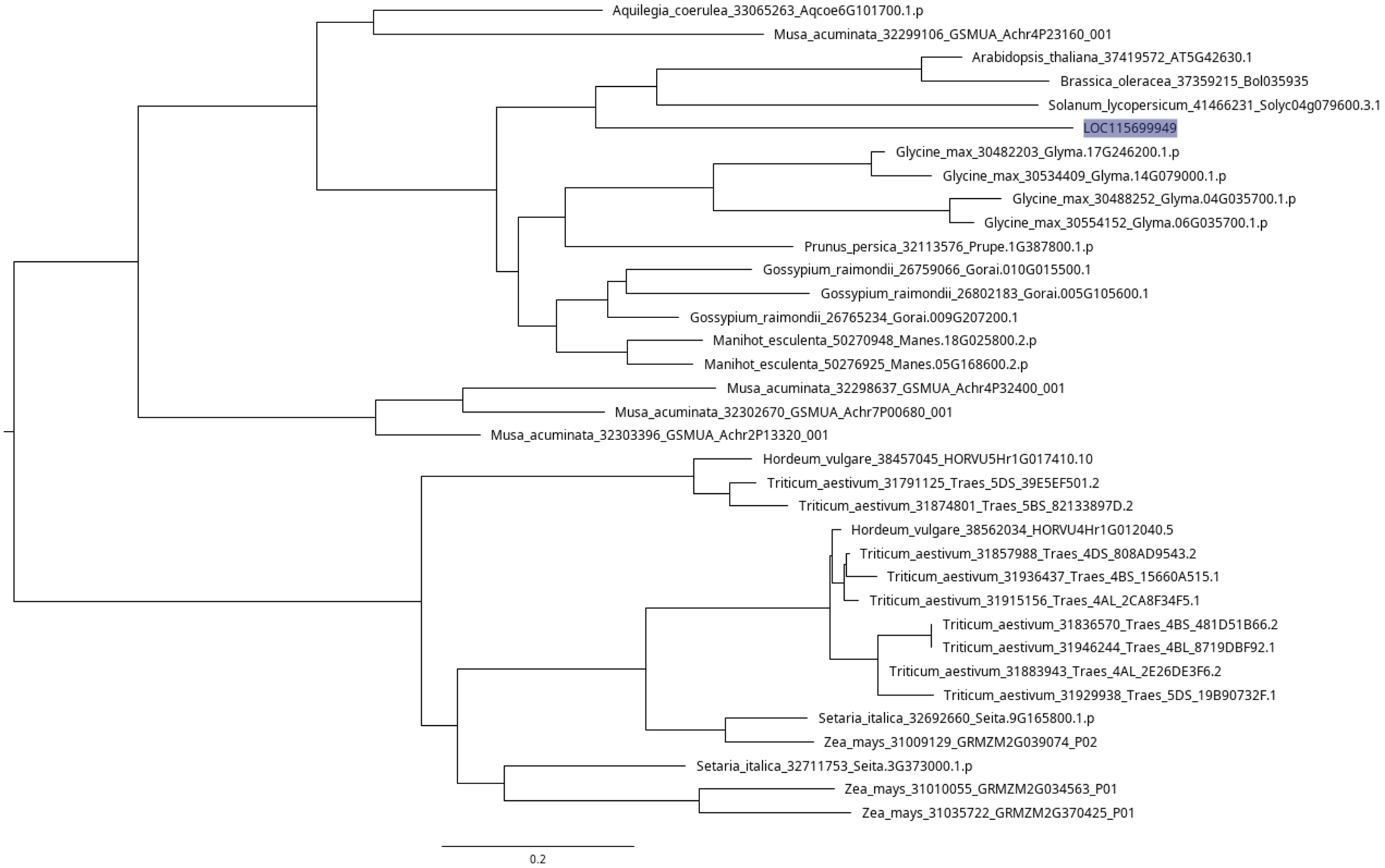


**Figure S13. Phylogenetic reconstruction of KAN4/ATS homologs.**
Maximum-likelihood phylogeny of KANADI family proteins from shoot.bio with *CsKAN4* (LOC115699949) highlighted. The tree demonstrates orthology between *Cannabis KAN4* and *Arabidopsis KANADI4/ABERRANT TESTA SHAPE* (*ATS*) (AT5G42630).

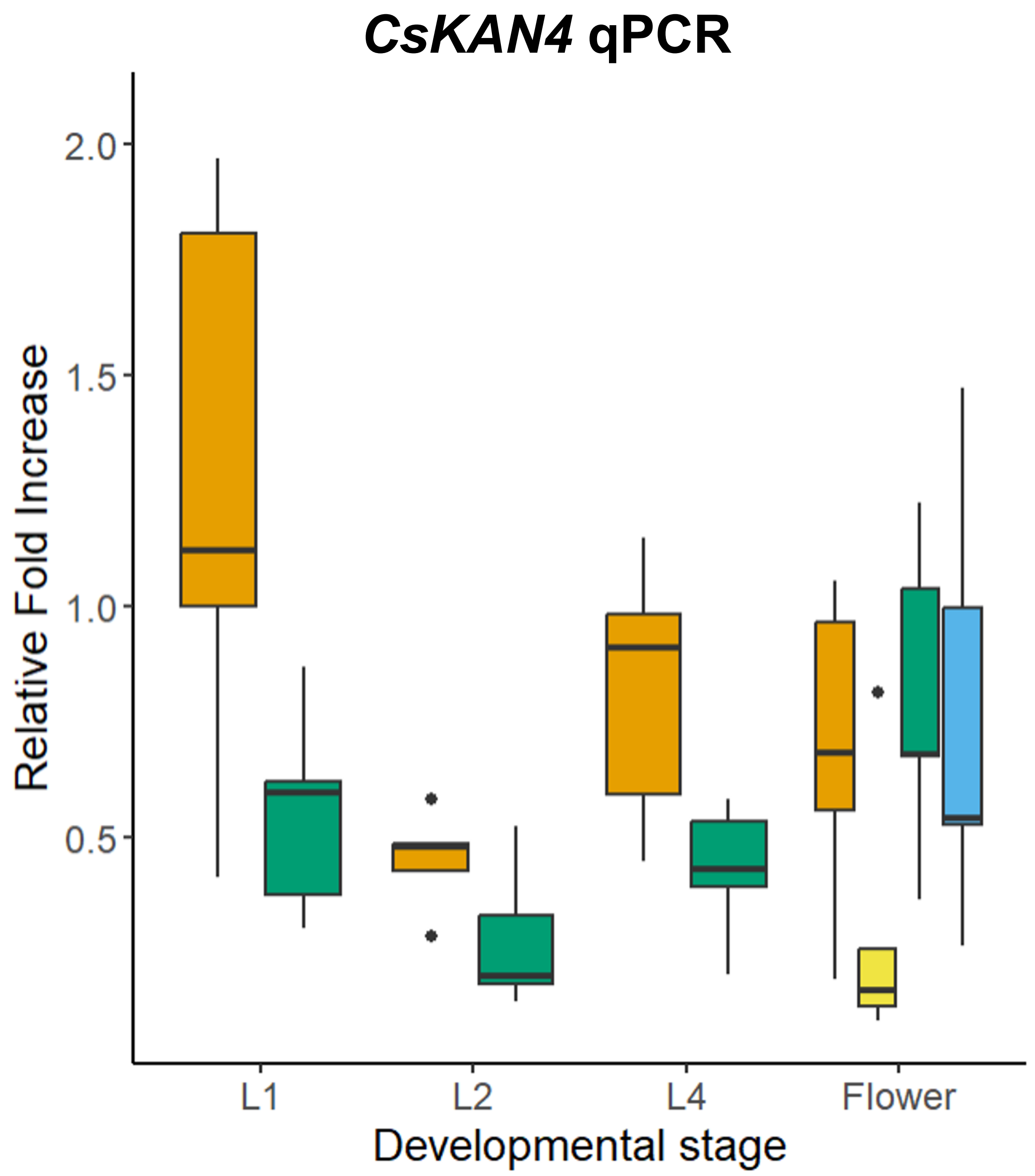


**Figure S14.** ***CsKAN4*** **qPCR using alternative housekeeping gene**
qPCR showing CsKAN4 (LOC115699949) expression, normalized to the *CsUBQ* housekeeping gene, in the apical meristem in early stages of plant development (two leaf pairs, stage L2) and in later stages ( four leaf pairs L9) and in flower tissue in cultivars ‘Felina 32’ (monoecious) and ‘FINOLA’ (dioecious).

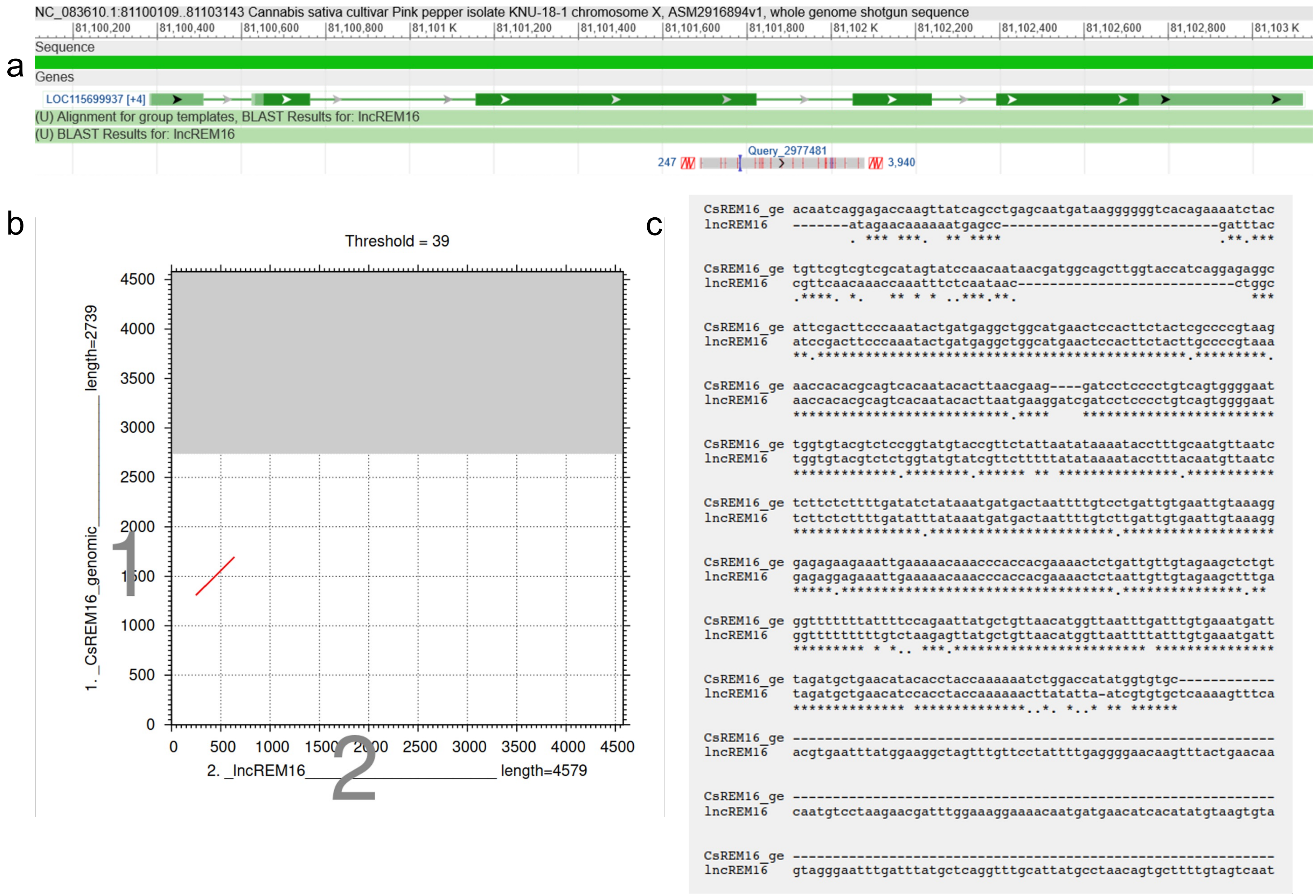


**Figure S15. Sequence similarity between *lncREM16* and *CsREM16*** a) Alignment from blast showing that a section of lncREM16 aligns with the genomic region of CsREM16 spanning part of the third and forth exons and the entire intron between the two exons b) dotplot and c) alignment from mafft of the matching regions.

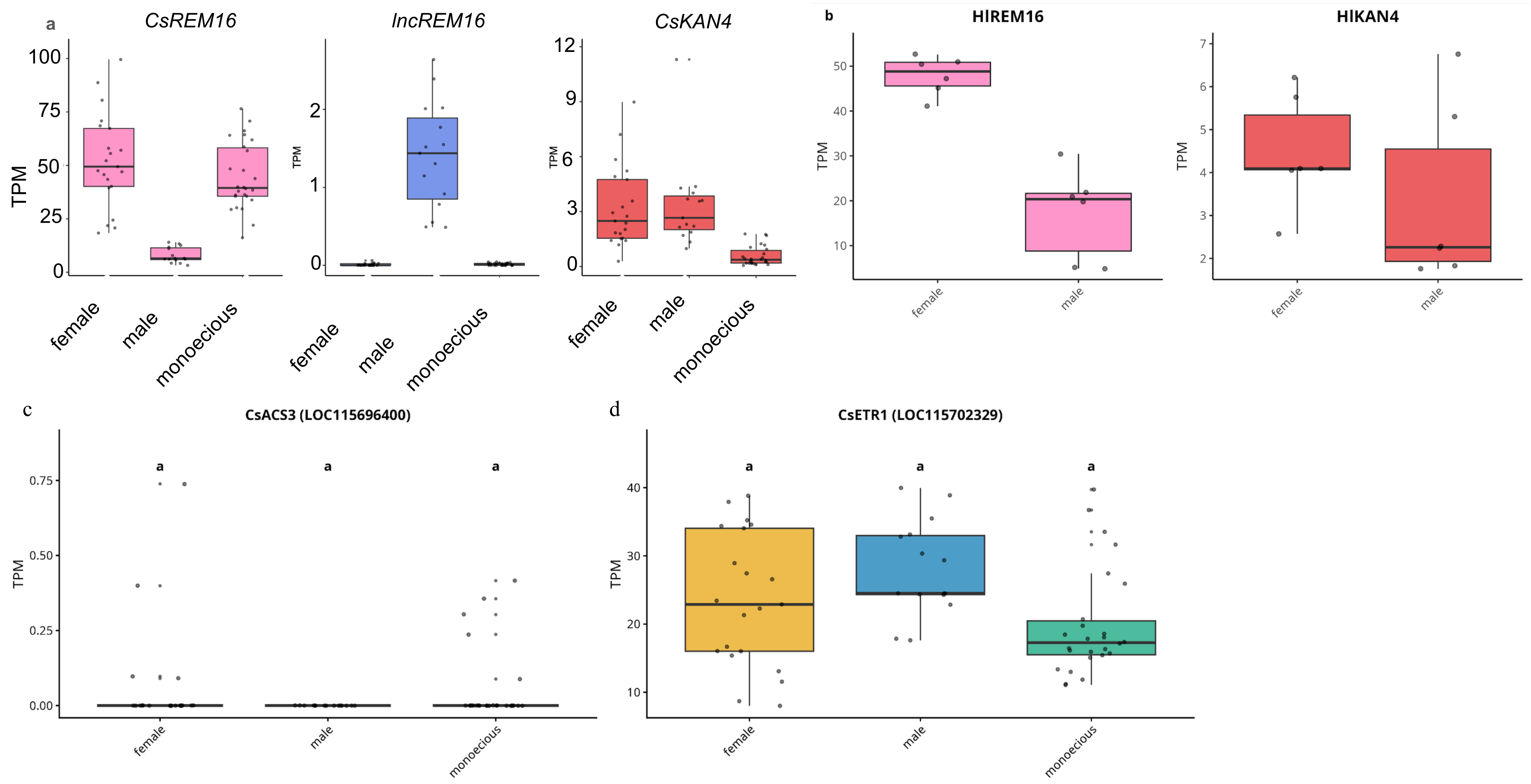


**Figure S16. Gene expression of the candidate sex determining genes**

**a)** Expression analysis of the three key genes using RNA-seq data from 64 'Felina 32', 'FINOLA' and F2 samples across different sex phenotypes. Box plots show transcript abundance (TPM) for *CsREM16* (LOC115699937), *lncREM16* (LOC133032448), and *CsKAN4* (LOC115699949) in female, male, and monoecious individuals. *CsREM16* shows high expression in females and monoecious plants but low expression in males, supporting a female-determining role. lncREM16 exhibits male-specific expression despite being X-linked. *CsKAN4* displays reduced expression in monoecious plants compared to dioecious individuals, consistent with its proposed role in monoecy determination. Individual data points are overlaid as jitter plots. **b)** Expression analysis of *H. lupulus* orthologous genes using RNA-seq data from five male and five female plants (BioProject PRJNA694508). *HlREM16* exhibits significant female-biased expression ($p < 0.05$), mirroring the expression pattern of *CsREM16* in Cannabis. *HlKAN4* shows similar expression levels between sexes. **c)** *CsACS3* expression is consistently near zero across all samples. **d)** *CsETR1* shows no significant expression differences between sexes or between dioecious and monoecious cultivars (adjusted p-value > 0.05). Box plots show transcript abundance (TPM) with median (center line), interquartile range (box), and 1.5× interquartile range (whiskers). Individual data points are overlaid.

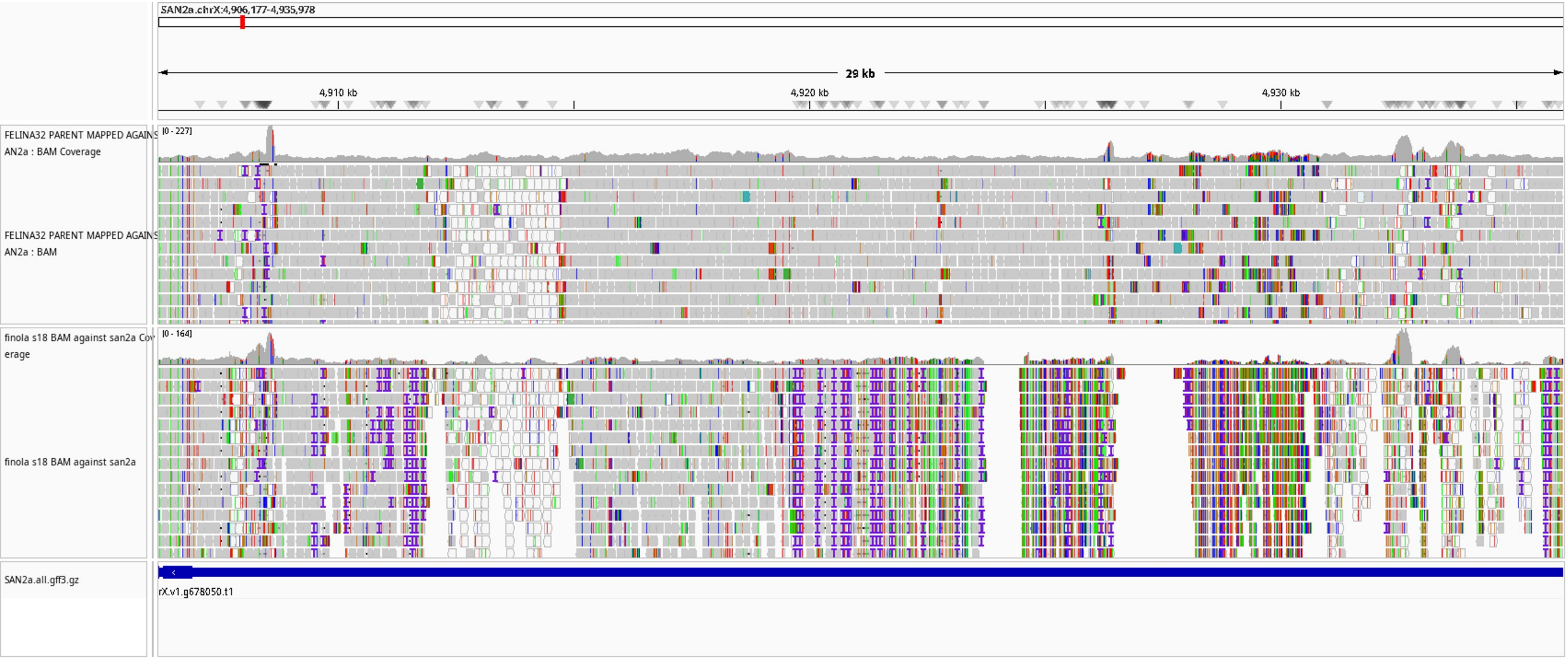


**Figure S17. Transposable element presence in monoecious cultivars Santhica27 and Felina 32.**
A transposable element (sequence in Supplementary Data 1) was identified upstream of the transcription factor *CsKAN4* in the monoecious cultivar 'Santhica27'. The presence of this element was confirmed in the monoecious cultivar 'Felina 32' by mapping sequencing reads from the 'Felina32' parent to the 'Santhica27' assembly and verifying coverage in this region. The absence of mapped reads from the dioecious cultivar 'FINOLA' in this region indicates the transposable element is absent in ‘FINOLA’.

a

```
HlREM16     MGDSCMECTKWAEEIYWTHFQDLHFSQIMATGFDRQLAIPKKFSSSLRNKLPEFVTLKGP
CsREM16     MGDSCVECTKWAEEIYWTRFQDIHFSQIMASGFDRHLAIPKKFSSSLREKLPECVTLKGP
            *****:************:***:*******:****:************:**** ******

HlREM16     GGGTWSVGVTTTGDTLYFKHGWQEFVNDHGLVENDLLIFKYSGKSHFEVLIFNGQNMCEK
CsREM16     GGGTWSVGVTTTGDTLYFKHGWQEFVNDHGLAENDLLIFKYSGKSVFEVLIFNGQNMCEK
            *******************************.************* **************

HlREM16     EFSYFVKRCGHKKDDHGGSLIKKKSRRASSDKSNNGSFENVECTSVEKSRNFDGLWVQSG
CsREM16     EFSYFVKICGHKKDDH-GTLSKKVSRRTSCDETNNDSFENVECTQVKKSRNFDGLWLQSG
            ******* ******** *:* ** ***:*.*::**.********.*:*********:***

HlREM16     EPVITLTNDQVVRRRS-VNRHIVSSSNDGSLVPSGEPFITPTTDEQVGINATSFRPIRTT
CsREM16     DQVISLSNDKGGHRKSTVRRRIVSNNNDGSLVPSGEAFDFPNTDE-AGMNSTSTRPVRTT
            : **:*:**:  :*:* *.*:***..**********.*  *.*** .*:*:** **:***

HlREM16     RNHNAINEESSPGSGELLYVSEAEHTPTKRSGASYGVQFLSNRRPVSEDEKNYALKLAHS
CsREM16     RSHNTLNEGSSPVSGELVYVSDAEHTPTKKSG-PYGVQYSSNRRPVTEDEKNFALQLAHS
            *.**::** *** ****:***:*******:** .****: ******:*****:**:****

HlREM16     EILESNSEGLIVVMKPSHVYKRFFVLLPTDWVMKYISLENQDVFLRFGEKEWRTRFNYNQ
CsREM16     QILESNSEGLIVVMKPSHVYKRFFVLLPTDWVMKYISLENQDVFLRYGEREWRTRFNYNQ
            :*********************************************:**:**********

HlREM16     LRKSAALSSGWNHFAVDNNLEEFDVCVFQPGSPVNNSFCLDVKIFRVVEEITPLTTISSP
CsREM16     LRKSAALSSGWNHFATDNNLEEYDVCVFQPGSPVNDSFCLDVKIFRVVEEITPLTTISSS
            ***************.******:************:***********************.

HlREM16     ISRKNKRKLIKNEKQSIV
CsREM16     -SRKYKRKLIKMDGQSIV
             *** ****** : ****
```

b

```
HlKAN4      MFPNQSTKM-------------------RSSSSSLPDLSLQISPPSIP---VPDCLTEAN
CsKAN4      MFPNHTTKMMRRTSITTATTSSSSSSSSSSSSSSMPDLSLQISPPSIPDYYHAITTTEAN
            ****::***                    *****:*************    .   ****

HlKAN4      N-----NEVVLLLSDRSSTTTDSGSS-TTGSDLSHENGLYNNNPLQKTTNCYSSLGGGSN
CsKAN4      NNKNNSNEVLLL---RSSTTTDSGSSTTTGSDLSHENGLEIKTTI--TTNCYN-------
            *     ***:**   *********** ************  :..:  *****.       

HlKAN4      HHEPTLSLGFETKDNHLPPP-------VLLHHQHHHHLPR-SFNSNSNNININ-NHHHFH
CsKAN4      -QEPTLSLGFETKDNNPPPPPVSVVPAVLL-----HHLPRSSFNNTSTNNNISIDHHKYH
             :*************: ***       ***     ***** ***..*.* **. :**::*

HlKAN4      SH-HQPQIYGRE---FKRNPRSISGV-KRSVRAPRMRWTTTLHAHFVHAVQLLGGHERAT
CsKAN4      FHGHQAQIYGRSGELMKRNPRSIGGVMKRSVRAPRMRWTTTLHAHFVHAVQLLGGHERAT
             * **.*****.   :*******.** *********************************

HlKAN4      PKSVLELMNVKDLTLAHVKSHLQMYRTVKSTDKAA--GQEQTDLGLNQ---------RTG
CsKAN4      PKSVLELMNVKDLTLAHVKSHLQMYRTVKSTDKAAGQGQGQTDLGLNQIRRVVGINNNNN
            ***********************************  ** ********         ...

HlKAN4      INIVNDDH-----VDGVLSSEILAQPHSGLPSSHVGPWSSLPVESPNERSHSSHENGSTV
CsKAN4      KNIVNDEHEHVDGSDGVLSSEILAQPHLGLPTNSHSHVSTRPI----ERSISSHENGWTV
             *****:*      ************* ***:.  .  *: *:    *** ****** **

HlKAN4      P---HDFGENGAKVYVEKKRSDSLEERVECSSLSSSNMCLNLEFTLGRPSWQKDYA-HHD
CsKAN4      ADHDQDFRENGAK--VEKVSSGS-----SLSLSSSSNMRLNLEFTLGRPSWQKDYAHHHD
            .   :** *****  ***  *.*     . *  ***** ***************** ***

HlKAN4      SSNE-LTLLKC
CsKAN4      SSNEVLTLLKC
            **** ******
```

**Figure S18. Protein sequence conservation between *CsREM16* (LOC115699937) / *HlREM16* (LOC133803719) and between *CsKAN4* / *HlKAN4* (LOC133804030)**

**a)** Pairwise amino acid sequence alignment of CsREM16 (LOC115699937) and HlREM16 (LOC133803719), showing 83.3% identity.

**b)** Pairwise amino acid sequence alignment of CsKAN4 and HlKAN4 (LOC133804030), showing 59.3% identity. Alignments were performed using [MAFFT/Clustal Omega]. Symbols denote fully conserved residues (*), amino acids with strongly similar properties (:), and with weakly similar properties (.).

a

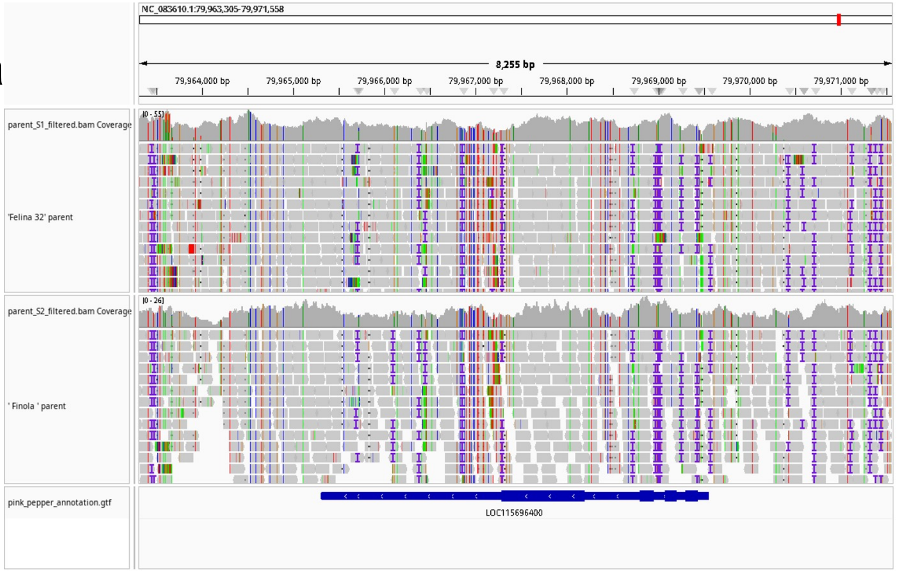


b

```
CLUSTAL format alignment by MAFFT (v7.511)


felina      MSSLLSRKAACNAHGQDSSYFLGWEEYERNSYDRITNPEGIIQMGLAENQLCYHLLESWL
finola      MSSLLSRKAACNAHGQDSSYFLGWEEYERNSYDRITNPEGIIQMGLAENQLCYHLLESWL
            ************************************************************

felina      ANNPHALGFRSQGQYIFRKLALFQDYHGLPEFKKAMVEFMSEISGNKVRFEPQSLVLIAG
finola      ANNPHALGFRSQGQYIFRKLALFQDYHGLPEFKKAMVEFMSEISGNKVRFEPQSLVLIAG
            ************************************************************

felina      ATSANEALIFCLADPNDAFLLPTPYYPGFDRDLKWRTEVVIVPIHCKSSNGFQITEEGLE
finola      ATSANEALIFCLADPNDAFLLPTPYYPGFDRDLKWRTEVVIVPIHCKSSNGFQITEEGLE
            ************************************************************

felina      QAYEDATNRNLRVKGVLITNPSNPCGTTMTVDELNLLLNFIELKKIHLISDEIYSGTVFN
finola      QAYEDATNRNLRVKGVLITNPSNPCGTTMTVDELNLLLNFIELKKIHLISDEIYSGTVFN
            ************************************************************

felina      KPDFKSVIEVLNERNNNNNNSTNQIMIREQVHVVYSLSKDLGLPGFRVGAIYSNHKMVLD
finola      KPDFKSVIEVLNERNNNNNNSTNQIMIREQVHVVYSLSKDLGLPGFRVGAIYSNHKMVLD
            ************************************************************

felina      AATKMSSFGLVSSQTQFLLSVMLSDKYFTKTYIKENQKRLKRRHKMLVGGLKKAGISCLK
finola      AATKMSSFGLVSSQTQFLLSVMLSDKYFTKTYIKENQKRLKRRHKMLVGGLKKAGISCLK
            ************************************************************

felina      SNAGLFCWVDMRHLLKSNTFNAEIELWKKIIYDVKLNISPGSSCHCNEPGWFRVCFANMS
finola      SNAGLFCWVDMRHLLKSNTFNAEIELWKKIIYDVKLNISPGSSCHCNEPGWFRVCFANMS
            ************************************************************

felina      GRTLKLAIKSVIESSLPLN
finola      GRTLKLAIKSVIESSLPLN
            *******************
```

**Figure S19. No sequence divergence was found between the monoecious and dioecious versions of the ethylene biosynthesis gene *CsACS3* LOC115696400.**

**a)** Sequencing reads from the monoecious 'Felina 32' and dioecious 'FINOLA' parents aligned to the reference Pink Pepper genome across the *CsACS3* locus. Coverage tracks show uniform read depth across both cultivars *with* no observable polymorphisms in coding regions. **b)** Protein sequence alignment of *CsACS3* translated from consensus sequences derived from genomic reads of 'Felina 32' and 'FINOLA' parents. The alignment reveals complete amino acid sequence identity between the monoecious and dioecious cultivars, indicating that *CsACS3* coding sequence is conserved.

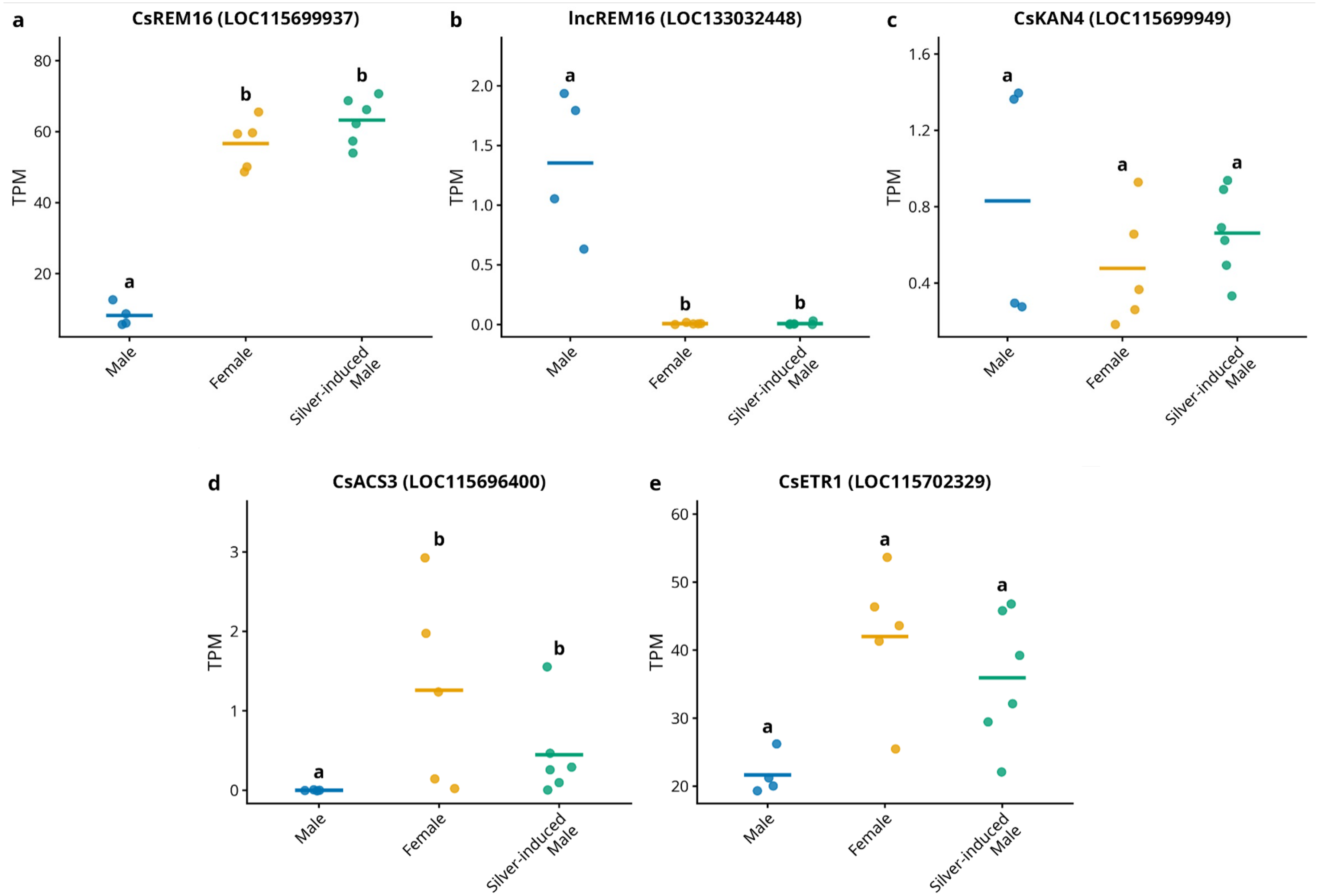


**Figure S20 Expression of the sex determination candidate gene in male, female, and silver-induced male flowers.**
**a)***CsREM16* (LOC115699937) shows high expression in female and silver thiosulfate-induced male (XX) flowers, with markedly lower expression in male (XY) flowers, indicating that expression follows genotypic sex (XX vs XY) rather than phenotypic sex.**b)** *lncREM16* (LOC133032448) displays male-specific expression with no detectable expression (TPM = 0) in female or silver-induced male flowers, consistent with Y-linked regulation. **c)** *CsKAN4* (LOC115699949) shows low expression across all three groups, consistent with reduced expression at the late flowering stage. **d)** *CsACS3* (LOC115696400) expression is silenced in male flowers, and is not significantly differnet between female and silver-induced male flowers. **e)** *CsETR1* (LOC115702329) shows higher expression in female and silver-induced male flowers compared to male flowers. Blue: male (XY); orange: female (XX); green: silver thiosulfate-induced male (XX). RNA-seq data from Adal et al. (2021), BioProject PRJNA669389.
