## Supplementary Text for "An ancient X chromosomal region harbours three genes potentially controlling sex determination in *Cannabis sativa*"

**Supplementary Text 1: Analysis of unexpected X chromosome genotypes in F2 population**

**Genotype expectations vs genotype observations**

To generate our mapping population, we used a ‘FINOLA’ male pollen donor to pollinate a ‘Felina 32’ plant. F1 males will therefore possess a ‘Felina 32’ X chromosome (termed here $X_M$) and a FINOLA Y chromosome, while F1 females inherit one X chromosome from the ‘FINOLA’ parent and one from the ‘Felina 32’ parent (F1 female genotype $X_DX_M$). These expected F1 genotypes were confirmed through whole genome sequencing. Consequently, $F_2$ female and monoecious plants derived from crossing $F_1$ individuals should theoretically possess either heterozygous ($X_DX_M$) or homozygous Felina 32 ($X_MX_M$) genotypes at X-linked markers, but not homozygous FINOLA ($X_DX_D$) genotypes:

| | | | | |
|---|---|---|---|---|
| P | $X_MX_M$ | | ⊠ | $X_DY$ |
| $F_1$ | $X_MX_D$ | | ⊠ | $X_MY$ |
| F2 | $X_MX_M$ | $X_DX_M$ | $X_MY$ | $X_DY$ |

Contrary to this expectation, approximately 20% of $F_2$ female and monoecious plants displayed $X_DX_D$ genotypes at X chromosome markers.

Noteworthy, recombination can take place in $F_1$ $X_MX_D$ individuals, so X chromosomes of $F_2$ plants can be a mix of $X_M$ and $X_D$ markers due to crossovers. Detailed analysis revealed that in 26 $F_2$ individuals (13% of the XX population) > of 80 % of the markers on the X chromosomes were homozygous $X_DX_D$. These individuals were non-randomly distributed among F1 families: Offspring of the F1-093 mother produced 16 of these 26 plants, while the F1-091 mother produced only one.

This skewed distribution correlates with sex ratio distortion, as F1 families producing more $X_DX_D$ genotype individuals also showed the greatest deviation from the expected 1:1 male:female ratios. Overall, only 20.1 % of the F2 plants were male, less than the expected

50% ($\chi^2$ test, $p < 0.001$). But while F1-093 mother produced only 15.9 % males, the F1-091 mother produced 35.9 % male plants (Supplementary Table 1).

The genetic data outlined above would be in agreement with some $F_1$ plants being selfed: as $F_1$ plants are $X_DX_M$, selfing could explain the lack of males as well as the occurrence of $X_DX_D$ genotypes in the $F_2$. However, no male flowers were observed on female $F_1$ plants and even if a small number of male flowers did develop it's hard to imagine they pollinated such a large number of female flowers that it can explain the large number of $X_DX_D$ genotypes we observed. An alternative explanation for the observed genotypes is automixis (not to be confused with apomixis), for example the fusion of two haploid megaspores. Though automixis is known from animals, reports from plants are very rare (https://link.springer.com/article/10.1007/s00497-024-00499-6). It will be interesting to see whether our observations are due to automixis in *C. sativa.*

The genetic details of the skewed genotype distribution of our $F_2$ population notwithstanding, it is important to note that the QTL analysis methodology performed here relies on the statistical associations between marker genotype frequencies and phenotypes within the $F_2$ population, regardless of the underlying mechanism that generated these genotypes. Whether the unexpected $X_DX_D$ genotypes arose from self-pollination of $F_1$ females or other biological mechanisms, the mathematical framework of QTL detection that identifies genomic regions at which genotype frequencies correlate with phenotypic variation stands.

**QTL validation by constraining the data before the analysis**

To further ensure these unexpected genotypes did not artificially create the observed QTL, we repeated the analysis using the R/qtl's sex-specific inheritance model. This model incorporates expected $F_1$ genotypes (heterozygous female × hemizygous male), and it automatically flags markers showing significant segregation distortion from expected Mendelian ratios, and identifies theoretically impossible genotypes based on the specified $F_2$ cross type. After setting all the unexpected $X_DX_D$ genotypes and the ones on the X chromosome with significant distortion compared to the one that was theoretically expected, to missing values, the QTL analysis maintained the exact same peak location on the X chromosome, with the most significant marker remaining the same, albeit with reduced LOD scores (Supplementary Fig. S5, S6). This validation confirms that while the origin of $X_DX_D$ genotypes remains unclear, their presence does not compromise the identification of the *Monoecy1* locus.

### Supplementary Text 2: Male-specific *lncREM16*

The long non-coding RNA *lncREM16* that we identified in our putative Sex Determining Region has exclusive expression in male plants and complete absence of expression in female plants. Based on the NCBI RefSeq annotation, *lncREM16* is annotated as gene ID LOC133032448, located on chromosome X at position 81,117,413. Interestingly, our RNA-seq data indicate that transcription of *lncREM16* initiates approximately 4,000 bp upstream of the annotated transcription start site (TSS). This observation is supported by the presence of multiple RNA-seq reads spanning both this upstream region and the annotated body of *lncREM16*, suggesting that the current annotation may underestimate the full length of its transcribed region.

Intriguingly, it is this upstream region (absent from the RefSeq annotation but actively transcribed in our male samples) that shares high sequence identity with a portion of the *CsREM16* gene. Specifically, it aligns with two exons and the intervening intron of *CsREM16*, suggesting a potential regulatory interaction. Given that *CsREM16* is transcriptionally active in females but silent in males (where the *lncREM16* is expressed), this reciprocal expression pattern raises the possibility of a regulatory or silencing mechanism involving the *lncREM16* in male plants.

As illustrated in the figure below, RNA-seq reads from a representative male sample clearly span both the upstream region and the body of the *lncREM16*, whereas no reads are detected in the corresponding region of female samples, confirming its strict male specificity.

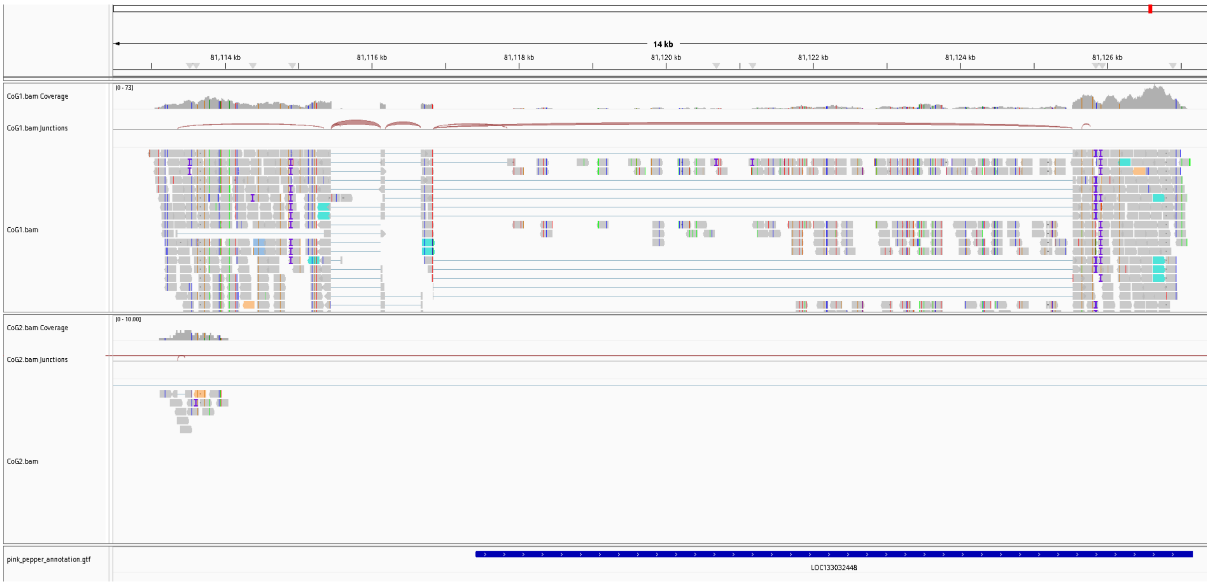
